# Postmortem tissue biomarkers of menopausal transition

**DOI:** 10.1101/2024.06.20.599941

**Authors:** Maria Tickerhoof, Heining Cham, Anaya Ger, Sonola Burrja, Pavan Auluck, Peter J. Schmidt, Stefano Marenco, Marija Kundakovic

**Author notes:** Correspondence should be addressed to M.K.

## Abstract

The menopausal transition (MT) is associated with an increased risk for many disorders including neurological and mental disorders. Brain imaging studies in living humans show changes in brain metabolism and structure that may contribute to the MT-associated brain disease risk. Although deficits in ovarian hormones have been implicated, cellular and molecular studies of the brain undergoing MT are currently lacking, mostly due to a difficulty in studying MT in postmortem human brain. To enable this research, we explored 39 candidate biomarkers for menopausal status in 42 pre-, peri-, and post-menopausal subjects across three postmortem tissues: blood, the hypothalamus, and pituitary gland. We identified thirteen significant and seven strongest menopausal biomarkers across the three tissues. Using these biomarkers, we generated multi-tissue and tissue-specific composite measures that allow the postmortem identification of the menopausal status across different age ranges, including the “perimenopausal”, 45-55-year-old group. Our findings enable the study of cellular and molecular mechanisms underlying increased neuropsychiatric risk during the MT, opening the path for hormone status-informed, precision medicine approach in women’s mental health.

## Introduction

The menopausal transition (MT) is the midlife transition period, typically lasting 2-8 years^1^, which extends from the point when menstrual periods become irregular (early MT) until the final menstrual period (FMP)^2^ (**Figure 1A**). Menstrual changes reflect changes in the function of the hypothalamic-pituitary-gonadal (HPG) axis (**Fig. 1B**) due to reproductive ovarian aging and declining ovarian follicle reserve. The entire MT is marked by the low antral follicle counts and low levels of the ovarian reserve marker anti-Müllerian hormone (AMH)^3,4^ (**Fig. 1B**). The levels of ovarian hormones estradiol and progesterone decrease, although this is also a period with the most extreme hormone fluctuations, particularly during the late MT^2^. To compensate for decreased ovarian function, the secretion of follicle stimulating hormone (FSH) from the pituitary (**Figure 1B**) is typically elevated and stays stably elevated postmenopause together with low ovarian hormone levels^5^. The hallmarks of MT and menopause are thus low/non-detectable AMH, low estradiol, and elevated FSH, and while cutoff values for blood AMH and FSH levels have been suggested^2,3^, these measures are not standardized as clinical menopausal biomarkers^2^.

**Figure 1.**
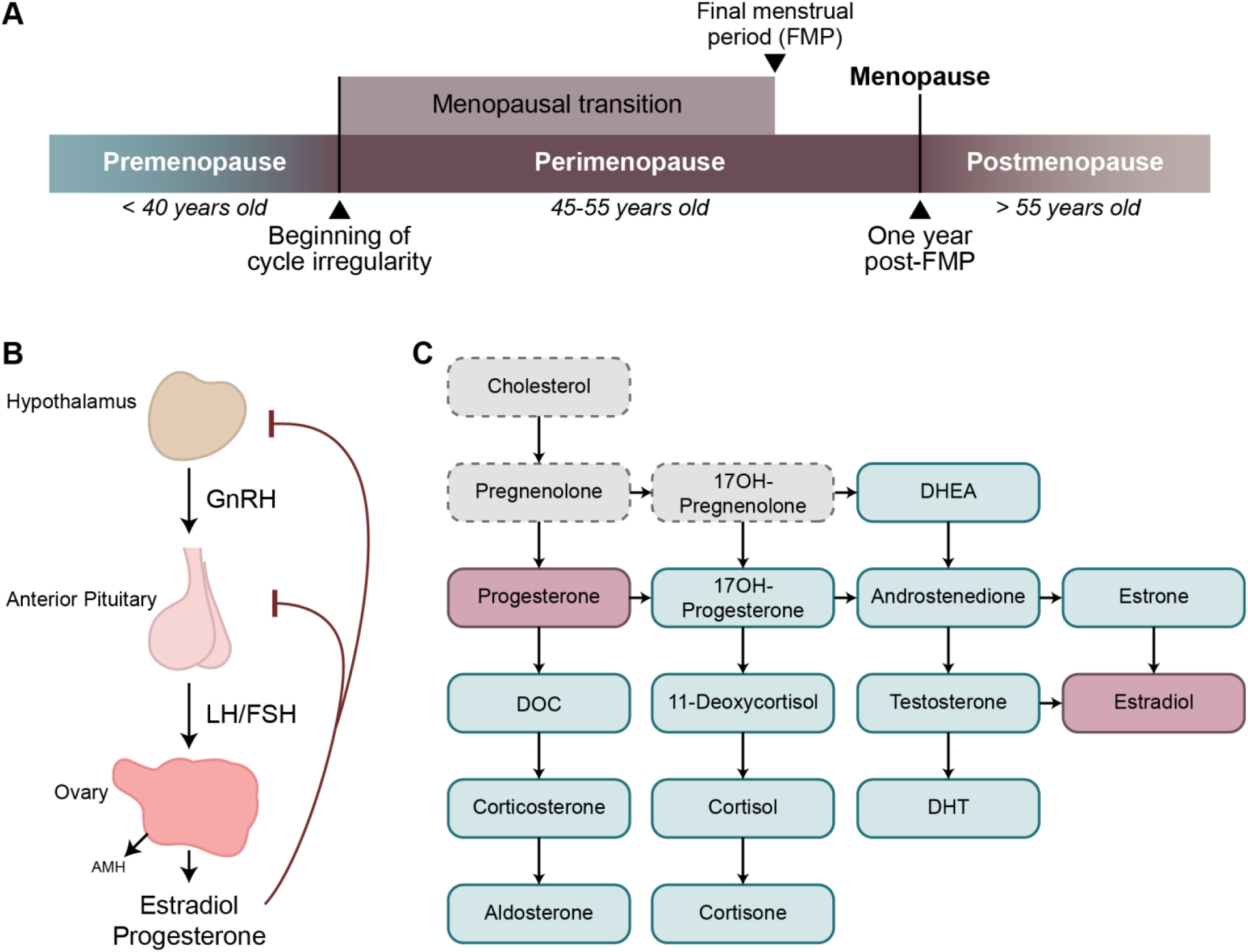
Characterizing menopausal transition using hypothalamic-pituitary-gonadal (HPG) axis-related markers: **A)** The menopausal transition is defined as the time between the beginning of cycle irregularity and the final menstrual period (FMP). One year after the FMP marks menopause, and following this point an individual is considered postmenopausal. Perimenopause includes the entire menopausal transition and one year post-FMP. **B)** Release of reproductive hormones is controlled by the HPG axis. GnRH is released from the hypothalamus into the hypophyseal portal system, signaling release of LH and FSH from the pituitary gland. From the pituitary gland, LH and FSH travel through the periphery until they reach the ovaries, which stimulates the release of estradiol and progesterone. Estradiol and progesterone provide feedback inhibition to the hypothalamus and pituitary gland to prevent release of GnRH, LH, and FSH. Within the ovary, AMH is produced by growing follicles, and serum AMH levels are an indicator of functional ovarian reserve. **C)** Fourteen of the seventeen key steroids involved in sex steroid synthesis were analyzed in the present study (blue; primary ovarian steroids in pink). Glossary: AMH – Anti-Müllerian hormone; DHEA – Dehydroepiandrosterone; DHT – Dihydrotestosterone; DOC – Deoxycorticosterone; FSH – Follicle-stimulating hormone; GnRH – Gonadotropin-releasing hormone; LH – Luteinizing hormone.

Clinically, menopause is diagnosed 12 months after the FMP, at the average age of 51 years^6^. The period of MT together with 12 months following the FMP is also known as *perimenopause (***Figure 1A**). Globally, >850 million women are aged 40–60 years and ∼90% of these women will transition through perimenopause within the age range of 40– 58 years^6^. The lack of physiologically meaningful, reliable, and objective criteria for classifying women into menopausal status categories has been impeding progress in understanding the effect of reproductive aging and MT on health and disease^7^.

While MT represents a natural life transition, it is associated with bothersome symptoms in up to 90% of women, including hot flashes, sleep disturbances, depressed mood, increased anxiety, and trouble concentrating^8^. Specifically, the MT is an important window of vulnerability for the development of mental disorders including mood and psychotic disorders. Studies showed a 2-5 fold increased risk for major depressive disorder (MDD) during the MT, compared with both late premenopause and several years post-menopause^9^. Similarly, women undergoing the MT exhibit a heightened risk for the first-onset psychosis and reemergence or exacerbation of symptoms of schizophrenia and related disorders, which are not observed in men of similar age^10^.

Although evidence suggests that hormonal changes play a critical role in the manifestation of mood and psychotic symptoms during the MT, the biological mechanisms are poorly understood^9-12^. Studies in living humans show changes in brain metabolism and brain structure that may contribute to the MT-associated psychiatric risk^13^. However, studies on cellular and molecular changes in the human brain during this period are currently lacking. Specifically, working with postmortem human brain tissue represents a particular challenge, as the major US brain banks do not typically have information regarding an individual’s reproductive status. While chronological age at death is informative, it does not represent a good proxy for the menopausal status, which can obscure the results of the investigations focused on the MT’s effects on the brain.

To enable cellular and molecular studies of MT in humans, here we characterize 39 candidate biomarkers in our cohort of 42 individuals across three postmortem tissues – blood, the hypothalamus, and pituitary gland. Our study identifies the strongest menopausal biomarkers across the three tissues and establishes a menopausal component score that allows for the postmortem determination of the menopausal status of any given individual, across the wide age range including the most challenging “perimenopausal” group, within 45-55 years of age. To make this analysis more cost-efficient, feasible, and adaptable for other researchers, we also offer alternative tissue-specific component scores for the reproductive status determination.

## Results

Here, we analyzed 39 candidate menopausal biomarkers including: i) a panel of 14 different steroid hormones (**Figure 1C**) in *blood and the hypothalamus*; ii) AMH in *blood*; iii) FSH protein levels in *blood* and *pituitary gland*; iv) gene expression of reproduction-relevant genes in the *hypothalamus* (*ESR1, ESR2, GPER, PGR, KISS1*) and *pituitary gland* (*FSH, ESR1, GNRHR*). Our cohort included 42 individuals whose postmortem tissue samples (blood, the hypothalamus, and pituitary gland) were received from the Human Brain Collection Core (HBCC) of the National Institute of Mental Health - Intramural Research Program. We originally classified the subjects based on their “expected menopausal status”, simply using their chronological age as follows: <40 years (“<u>pre</u>-menopausal”, N=10), 45-55 years (“peri-menopausal”, N=21), and >55 years (“<u>post</u>-menopausal”, N=11) (**Table 1**). While our target was the perimenopausal group, which is likely to show variability in the menopausal age of onset, we included younger (<40 years) and older (>55 years) groups to define pre- and post-menopausal reference values, respectively, for biomarkers. Notably, our cohort also included a significant number of individuals with MDD or depressive symptoms (N=18), in addition to individuals with anxiety symptoms or substance use (**Table 1**). Education level, post-mortem interval (PMI), and RNA integrity number (RIN) for the hypothalamus and pituitary gland were comparable across all three groups (**Table 1**).

**Table 1:** Subject basic information.

| Group | Pre-menopausal | Peri-menopausal | Post-menopausal |
| --- | --- | --- | --- |
| N | 10 | 21 | 11 |
| Age (years)* | 26 (17-39) | 49.6 (45-55) | 62.7 (56-67) |
| Education (years) | 15 (11-18) | 13 (10-18) | 14.5 (11-20) |
| Race (%Black/White/Other) | 40/60/0% | 29/62/9% | 36/64/0% |
| Hormonal contraceptive use | 40% | 14% | 0% |
| Hormone replacement therapy | 0% | 10% | 9% |
| PMI (hours) | 28.25 (14.5-42) | 28.5 (13-74.5) | 28 (14-44.2) |
| Hypothalamus RIN | 5.0 (2.6-5.8) | 5.1 (2.6-6.2) | 5.1 (2.7-6.2) |
| Pituitary RIN | 6.1 (5.3-6.7) | 5.8 (3.2-6.7) | 6.0 (5.2-6.7) |
| % Axis I confident | 30% | 67% | 45% |
| % MDD/Depression | 20% | 52% | 45% |
| % Anxiety | 10% | 0% | 18% |
| % Substance use | 30% | 43% | 0% |

Age, education, PMI, and RIN values are displayed as mean (range). There was a statistically significant difference in age between the groups (\*p < 0.05, one-way ANOVA), but no differences in other parameters. PMI, Post-mortem interval; RIN, the RNA integrity number.

### Biological measurements

A total of 39 analytes related to the HPG axis and steroidogenesis were measured in the blood,hypothalamus, and pituitary glands of the 42 samples (**Table 2**). To identify strongest menopausal biomarker candidates, we initially did statistical analysis on the three groups based on age: pre-, peri, and post-menopausal groups (**Fig. 1A**). Of the 39 markers, 13 had significantly different measurements between groups determined by non-parametric Kruskal-Wallis test. All 13 of these significant effects were driven by differences between the pre- and post-menopausal groups as determined by posthoc Dunn’s test (**Table 2**). Perhaps not surprisingly, in *blood*, we found significant differences between pre- and post-menopausal groups in AMH (p < 0.001), FSH (p < 0.001), estrone (p = 0.009), estradiol (p < 0.001), progesterone (p = 0.011), and DHT (p = 0.021). In *the pituitary gland*, three markers showed significant changes from pre- to post-menopausal groups including FSH protein levels (p = 0.002) and gene expression levels of *FSH* (p < 0.001) and *GNRHR* (p = 0.049). Finally, we found differences in the levels of four *hypothalamic* steroids between pre- and post-menopausal groups including DHEA (p = 0.005), estrone (p = 0.003), estradiol (p = 0.023), and progesterone (p = 0.042).

**Table 2:** Group values of all 39 measures. Highlighted rows indicate measures with a significant difference between groups (Kruskal-Wallis, p < 0.05), with all 13 markers’ significance being driven by differences between premenopause and postmenopause groups (Dunn’s test, p < 0.05).

|  | Premenopausal –<br>Median (IQR) | Perimenopausal –<br>Median (IQR) | Postmenopausal –<br>Median (IQR) | H(2) | p<br>(Kruskal-Wallis) | Pre vs. Post p<br>(Dunn's test) |
| --- | --- | --- | --- | --- | --- | --- |
| **Glycoproteins** |  |  |  |  |  |  |
| *Blood* |  |  |  |  |  |  |
| AMH (pg/mL) | 1113.34 (677.07-2078.45) | 8.82 (5.52-25.83) | 3.77 (0.23-9.76) | 22.70 | 0.000012 | 0.000012 |
| FSH (mIU/mL) | 6.41 (3.02-7.63) | 19.58 (11.48-56.88) | 52.38 (37.91-97.81) | 17.81 | 0.00014 | 0.000084 |
| *Pituitary gland* |  |  |  |  |  |  |
| FSH (IU/mg) | 5.61 (4.01-9.43) | 21.85 (10.99-32.45) | 30.42 (21.00-36.40) | 12.07 | 0.0024 | 0.0019 |
| **Gene expression** |  |  |  |  |  |  |
| *Pituitary gland* |  |  |  |  |  |  |
| FSH (rel. expression) | 1.06 (0.56-1.75) | 3.41 (1.18-12.36) | 17.91 (13.63-26.70) | 16.36 | 0.00028 | 0.0002 |
| ESR1 (rel. expression) | 1.35 (0.70-1.77) | 1.37 (1.14-2.12) | 2.32 (1.13-2.74) | 2.88 | 0.24 |  |
| GNRHR (rel. expression) | 0.83 (0.48-1.90) | 1.26 (0.40-3.12) | 3.10 (2.24-4.19) | 6.19 | 0.045 | 0.049 |
| *Hypothalamus* |  |  |  |  |  |  |
| ESR1 (rel. expression) | 1.13 (0.81-1.28) | 1.10 (0.86-1.46) | 1.06 (0.60-1.51) | 0.31 | 0.86 |  |
| ESR2 (rel. expression) | 0.89 (0.74-1.20) | 1.42 (1.11-1.92) | 1.24 (0.50-1.45) | 5.39 | 0.067 |  |
| KISS1 (rel. expression) | 0.66 (0.39-2.21) | 1.12 (0.27-2.35) | 1.04 (0.64-11.72) | 0.69 | 0.71 |  |
| GPER (rel. expression) | 0.89 (0.73-1.48) | 0.86 (0.56-1.26) | 1.03 (0.68-1.59) | 0.58 | 0.75 |  |
| PGR (rel. expression) | 0.96 (0.73-1.24) | 1.37 (1.03-1.63) | 1.29 (0.96-1.81) | 2.34 | 0.31 |  |
| **Steroids** |  |  |  |  |  |  |
| *Blood* |  |  |  |  |  |  |
| Aldosterone (pg/mL) | 85.30 (41.02-176.97) | 94.53 (19.58-284.84) | 85.07 (19.75-151.09) | 0.37 | 0.83 |  |
| Androstenedione (pg/mL) | 610.13 (457.51-885.19) | 429.47 (235.64-871.63) | 384.45 (145.78-633.67) | 2.66 | 0.26 |  |
| Corticosterone (pg/mL) | 5144.79 (2545.65-6983.63) | 3442.73 (853.14-9726.20) | 5014.16 (2566.57-8408.12) | 1.38 | 0.50 |  |
| Cortisol (pg/mL) | 51731.36 (46492.11-74805.49) | 62930.20 (19910.22-166069.72) | 69105.31 (58189.20-93352.45) | 0.78 | 0.68 |  |
| Cortisone (pg/mL) | 6439.26 (6124.90-9978.50) | 7804.68 (4845.40-17361.96) | 10336.42 (7476.86-10671.95) | 0.32 | 0.85 |  |
| 11-deoxycortisol (pg/mL) | 248.06 (214.18-573.49) | 187.28 (124.27-849.73) | 342.03 (176.99-563.14) | 0.060 | 0.97 |  |
| DHEA (pg/mL) | 4611.79 (3292.17-9312.06) | 2040.58 (0.00-6776.87) | 0.00 (0.00-1959.97) | 5.52 | 0.063 |  |
| DHT (pg/mL) | 132.18 (109.69-234.22) | 33.00 (0.00-71.75) | 33.00 (30.00-33.00) | 8.33 | 0.016 | 0.021 |
| DOC (pg/mL) | 28.80 (22.71-91.74) | 35.28 (12.57-164.18) | 48.37 (14.10-76.66) | 0.30 | 0.86 |  |
| Estrone (pg/mL) | 156.89 (49.68-245.43) | 25.28 (17.33-83.75) | 20.40 (17.00-33.18) | 9.22 | 0.010 | 0.0088 |
| Estradiol (pg/mL) | 446.39 (243.68-849.39) | 65.87 (25.95-160.66) | 23.98 (12.29-52.42) | 14.07 | 0.00088 | 0.00057 |
| 17OH-progesterone (pg/mL) | 279.41 (186.51-355.63) | 308.56 (133.95-573.75) | 127.87 (54.73-327.46) | 4.56 | 0.10 |  |
| Progesterone (pg/mL) | 477.26 (87.34-1429.87) | 85.17 (44.03-242.00) | 61.34 (18.40-84.41) | 8.59 | 0.014 | 0.011 |
| Testosterone (pg/mL) | 478.79 (410.99-687.78) | 278.80 (206.28-483.78) | 327.20 (187.34-358.31) | 4.53 | 0.10 |  |
| *Hypothalamus* |  |  |  |  |  |  |
| Aldosterone (pg/g) | 144.94 (30.00-227.14) | 30.00 (0.00-234.52) | 166.00 (7.50-179.71) | 0.66 | 0.72 |  |
| Androstenedione (pg/g) | 999.76 (736.27-1430.89) | 406.47 (188.67-1263.65) | 357.15 (91.37-523.30) | 5.53 | 0.063 |  |
| Corticosterone (pg/g) | 8867.67 (3472.11-18187.80) | 3216.42 (600.00-20691.34) | 5226.10 (2686.39-9429.71) | 1.81 | 0.41 |  |
| Cortisol (pg/g) | 20433.88 (15166.19-37791.22) | 15865.83 (8906.07-101410.12) | 21081.38 (14417.18-69108.67) | 0.20 | 0.91 |  |
| Cortisone (pg/g) | 1834.45 (847.12-2941.47) | 1335.27 (600.00-4220.00) | 1330.79 (600.00-1697.93) | 1.39 | 0.50 |  |
| 11-deoxycortisol (pg/g) | 485.86 (148.83-926.77) | 534.81 (60.00-2987.06) | 375.34 (137.10-734.68) | 0.087 | 0.96 |  |
| DHEA (pg/g) | 25405.86 (11806.28-37697.34) | 3600.00 (3600.00-16709.99) | 0.00 (0.00-9000.00) | 10.05 | 0.0066 | 0.0051 |
| DHT (pg/g) | 180.00 (180.00-180.00) | 180.00 (180.00-180.00) | 180.00 (180.00-180.00) | 0.0029 | 1.00 |  |
| DOC (pg/g) | 146.48 (44.57-334.23) | 47.28 (18.00-472.30) | 108.48 (18.00-209.10) | 0.20 | 0.90 |  |
| Estrone (pg/g) | 371.62 (169.70-570.48) | 126.86 (89.58-324.00) | 30.00 (30.00-102.57) | 10.73 | 0.0047 | 0.0034 |
| Estradiol (pg/g) | 236.72 (73.57-328.72) | 30.00 (30.00-178.19) | 30.00 (30.00-30.00) | 7.22 | 0.027 | 0.023 |
| 17OH-progesterone (pg/g) | 1138.62 (648.20-2170.98) | 390.91 (180.00-2970.40) | 457.92 (180.00-544.06) | 1.65 | 0.44 |  |
| Progesterone (pg/g) | 882.31 (497.73-6396.58) | 434.69 (147.53-1278.97) | 139.68 (60.00-307.58) | 6.06 | 0.048 | 0.042 |
| Testosterone (pg/g) | 422.80 (273.02-873.55) | 281.15 (203.68-564.39) | 234.33 (152.93-365.10) | 2.03 | 0.36 |  |

### Steroid hormones

All three of the major ovarian hormones – estrone (**Figure 2A**), estradiol (**Figure 2B**), and progesterone (**Figure 2C**) – were significantly lower in the postmenopausal group than the premenopausal group in both the blood and hypothalamic tissue. Because of potential issues with accessing blood from different postmortem tissue/brain repositories, we additionally wished to gauge whether the levels of these steroids within the hypothalamus correlated highly with those within the blood. Estrone levels had a very high correlation between the two tissue types (**Figure 2A-iii**; r = 0.95, p < 0.001), and a moderate correlation was also observed for both estradiol (**Fig. 2B-iii**; r = 0.44, p = 0.007) and progesterone (**Fig. 2C-iii**; r = 0.44, p = 0.006). Moderate to very strong correlations were also observed between blood and hypothalamus measurements of 9 of the other 11 steroids analyzed (**Supp. Fig. 1**). The only two steroids that did not have a strong positive correlation were DHT and testosterone, which are likely explained by technical limitations; DHT levels within the hypothalamus were below the limit of quantitation for every sample, and 6 samples were excluded from testosterone measurement due to technical issues with sample processing. Therefore, quantifying steroid hormones extracted from hypothalamic brain tissue could represent a proxy for steroid hormone levels in the absence of access to blood.

**Figure 2:**
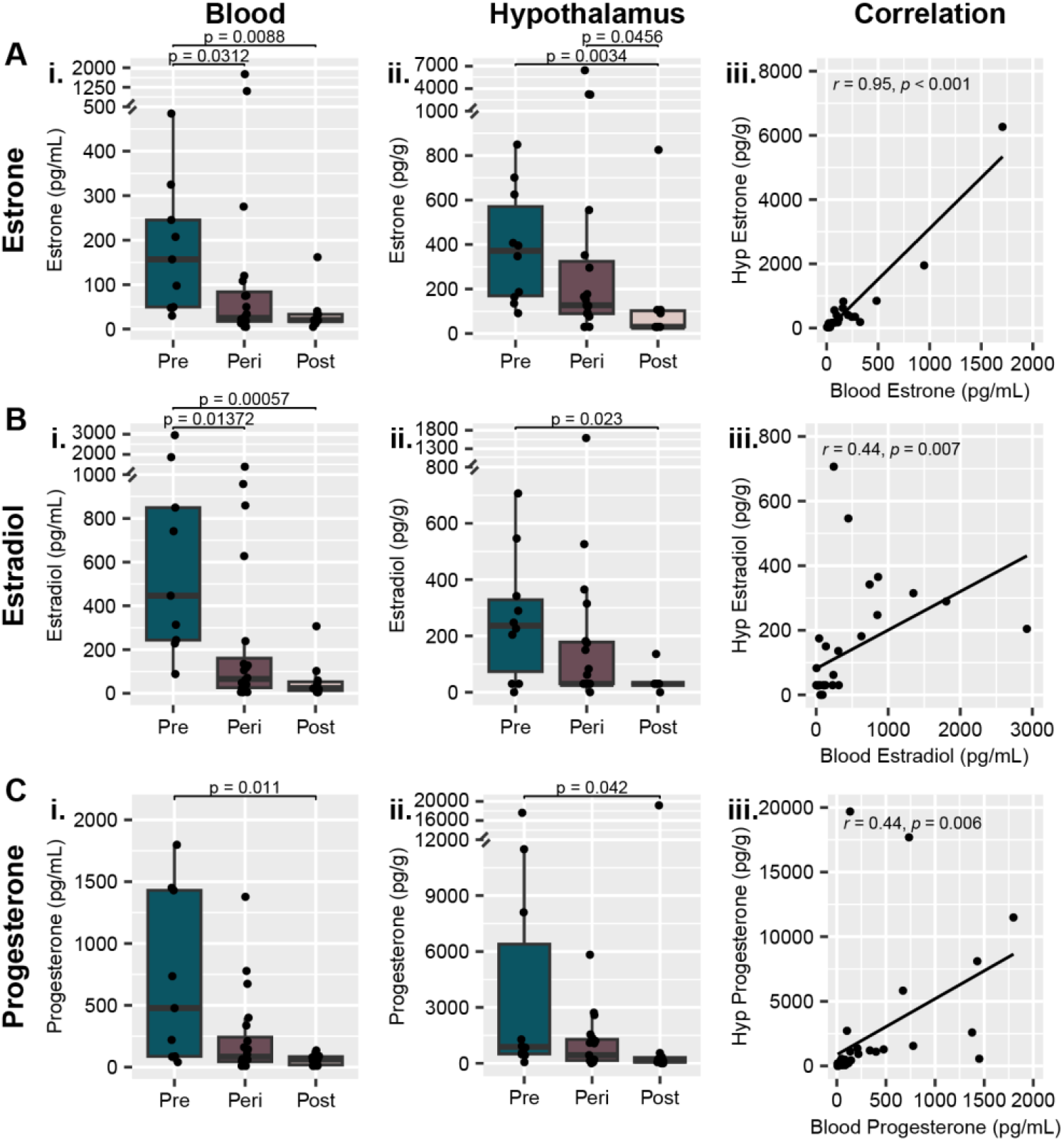
Ovarian steroids decrease from premenopause to postmenopause in both the blood and hypothalamus. **A)** Estrone, **B)** estradiol, and **C)** progesterone measurements are all lower in both **i)** blood and **ii)** hypothalamus tissue extracts. **iii)** Steroid measurements in the two tissue types had a high correlation for estrone and a moderate correlation for both estradiol and progesterone. Y-axis breaks indicate an adjustment in scale for display of individual value extremes. p values above box plots indicate significance of Dunn’s test posthoc comparisons following non-parametric Kruskal-Wallis test. Box plots (box, 1st-3rd quartile; horizontal line, median; whiskers, 1.5xIQR).

### Glycoproteins

AMH and FSH are often measured in clinical settings to assist with determining reproductive status^2^. Within our cohort of postmortem samples, there was a significant decrease in AMH (**Figure 3A**) from the premenopausal group to the postmenopausal group (p < 0.001). Notably, AMH was below 10 pg/mL in 8 of the 11 postmenopausal samples, indicative of a high likelihood of having reached the FMP^3^, with 5 samples having AMH levels below the limit of quantitation indicating a completely depleted ovarian reserve. Conversely, every premenopausal sample had AMH levels >100 pg/mL, indicating a very low chance that the FMP would occur within three years of the time of measurement^3^. Thus, our results align with expectations based on clinical observations in living humans^3^. Within the perimenopausal group, only 11 of the 20 samples had AMH levels below 10 pg/mL (three unquantifiable). The remaining nine measurements were all within the range of 10-100 pg/mL; these levels provide a low confidence that the FMP has occurred, as AMH may still be detected with residual ovarian function after the onset of amenorrhea, particularly within individuals younger than 51 years of age^3^. Thus, AMH measurements alone are not sufficient to characterize perimenopausal samples within our dataset.

**Figure 3:**
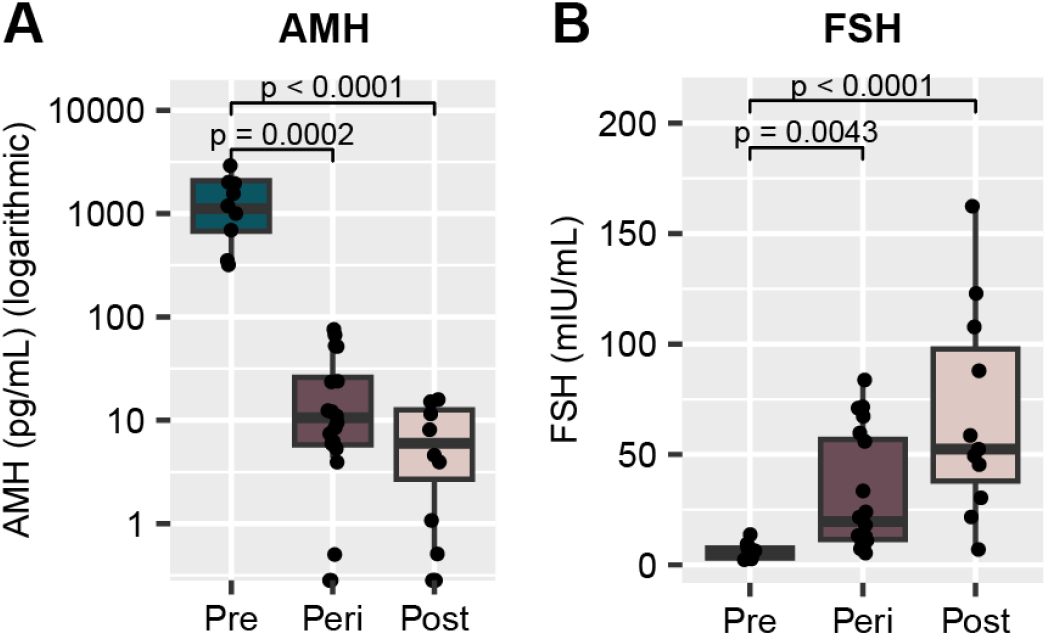
Glycoproteins related to ovarian function vary depending on reproductive status. **A)** AMH in the blood is significantly reduced from premenopause to perimenopause and postmenopause, and is nearly undetectable in the postmenopause group. AMH measurements are displayed on a logarithmic scale. **B)** FSH protein levels in the blood are significantly higher in the perimenopause and postmenopause groups when compared to premenopausal levels. p values above box plots indicate significance of Dunn’s test posthoc comparisons following non-parametric Kruskal-Wallis test. Box plots (box, 1st-3rd quartile; horizontal line, median; whiskers, 1.5xIQR); Glossary: AMH – Anti-Müllerian hormone; FSH – Follicle-stimulating hormone.

There was an inverse effect of MT on blood FSH levels, with a significant increase being observed between pre- and post-menopausal groups (p < 0.001, **Figure 3B**). Our measurements once again align with clinical observations^2^, with all premenopausal samples having measurements <15 mIU/mL and 9 out of 11 postmenopausal samples measuring >25 mIU/mL. Although FSH is often monitored in a clinical setting to help determine menopausal status, it alone is insufficient to diagnose menopause due to extreme variability during the MT, as well as susceptibility to factors such as hormonal contraceptives, hormone replacement therapy, and obesity^5,14^. Therefore, while aligning well with clinical observations, similar to AMH, FSH levels are insufficient to characterize the reproductive status of perimenopausal samples within our dataset.

### Pituitary gland

As a central component of the HPG axis, FSH production within the pituitary gland is likely significantly impacted across the menopausal transition, although this has not been addressed previously. Here we show that both FSH protein levels (p = 0.002, **Figure 4A**) and *FSH* gene expression (p < 0.001, **Figure 4B**) within the pituitary gland were significantly higher in the postmenopausal group compared to the premenopausal group, which aligns with the overall expected increase in FSH production and observed increase in blood FSH in the postmenopausal group. Additionally, expression of the gene for the GnRH receptor (*GNRHR*), which would trigger FSH production and release in response to GnRH released from the hypothalamus (**Figure 1B**), was also higher in the postmenopausal group (p = 0.049, **Figure 4C**). Once again, in the event that blood is not available to measure FSH, we wished to determine whether the pituitary gland could be an appropriate proxy for measuring FSH. Similar to the steroids, there was a significant moderate correlation between blood FSH and pituitary FSH (**Figure 4D**; r = 0.42, p = 0.007). Finally, we also confirmed that FSH protein and *FSH* expression within the pituitary had a significant moderate correlation (**Figure 4E**; r = 0.57, p < 0.001). This data shows that the pituitary gland can be used as an alternative tissue source for determining menopausal status *postmortem*.

**Figure 4:**
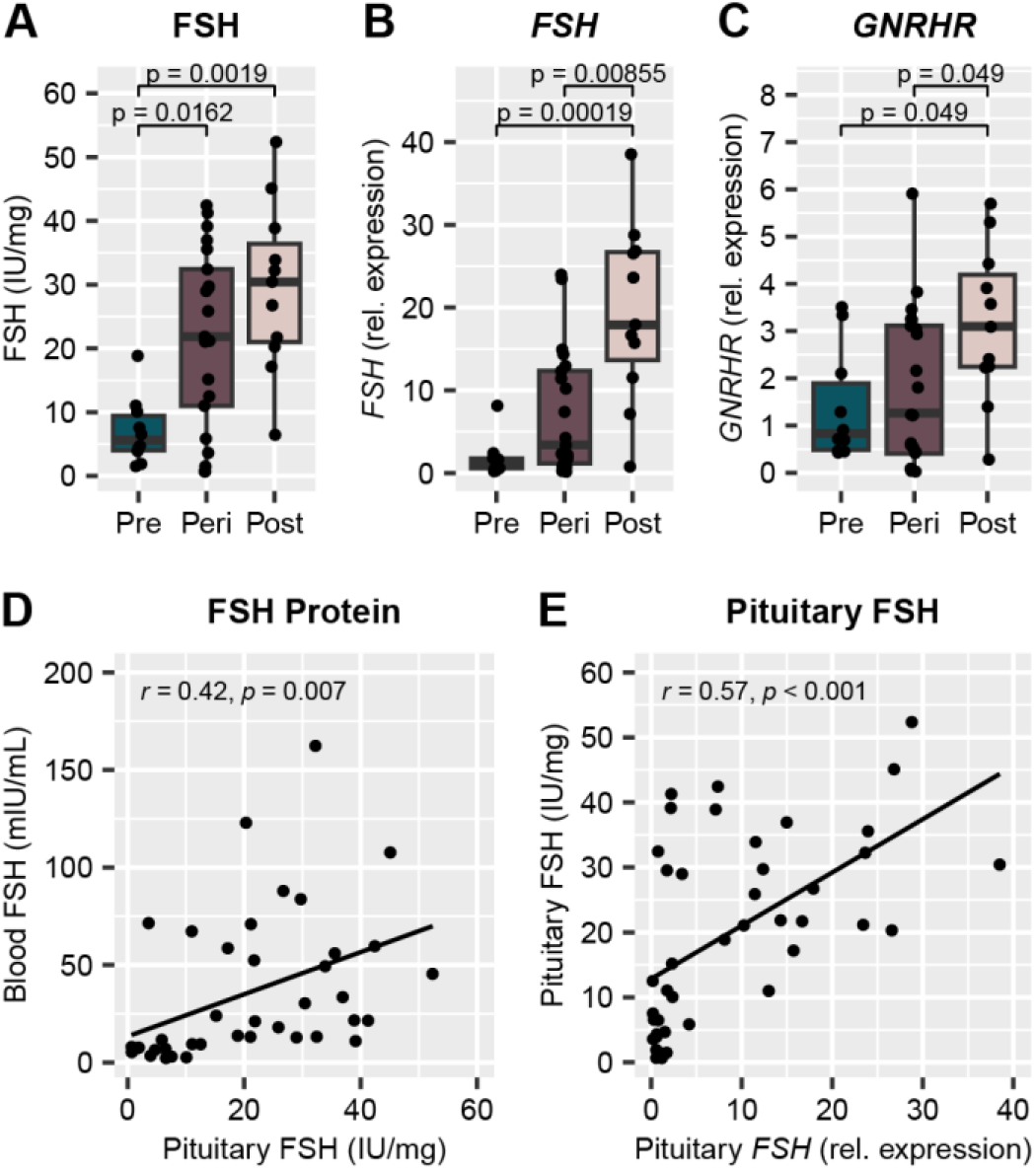
Pituitary gland FSH production increases across the menopausal transition. **A)** FSH protein levels and **B)** *FSH* gene expression within the pituitary gland are significantly increased in both the perimenopause and postmenopause groups when compared to the premenopause group. **C)** *GNRHR* gene expression within the pituitary gland is higher in the postmenopause group than the premenopause and perimenopause groups. There is a moderate correlation between **D)** blood FSH protein and pituitary gland FSH protein measurements, and **E)** FSH protein measurements and *FSH* gene expression within the pituitary gland. p values above box plots indicate significance of Dunn’s test posthoc comparisons following non-parametric Kruskal-Wallis test. Box plots (box, 1st-3rd quartile; horizontal line, median; whiskers, 1.5xIQR); Glossary: FSH – Follicle-stimulating hormone (protein); *FSH* – Follicle-stimulating hormone (gene); *GNRHR* – Gonadotropin-releasing hormone receptor (gene).

### Composite measurement

Although we discovered 13 biological measurements that significantly differed between the premenopausal and postmenopausal groups (**Table 2**), with the possible exception of AMH, each measure showed some overlap between pre- and post-menopausal individuals (**Figures 2-4**). Therefore, our next goal was to incorporate these markers into a single composite measure which we predict to be more informative than any of the single measurements. In this way, “perimenopausal samples” that could not be definitively categorized as pre- or post-menopausal based on age alone (no information whether the FMP had been reached was available) could have the composite score used to be characterized as statistically, biologically more similar to a “premenopause-like” or “postmenopause-like” group. In order to achieve this, principal component analysis (PCA) was performed to integrate biological measurements of interest into a single component score. PCA is a statistical procedure that identifies groups of variables according to degrees of association, and reduces each associated group into a new variable – so called “component.” Thus, this approach can be used to reduce correlating variables into a single component score to be used as our composite measure.

### Principal component analysis (PCA)

The first step of PCA was to examine the linearity and correlation between each of the 13 individual markers (**Supp. Figure 2**). Of the 78 comparisons drawn between each of the 13 markers, 34 comparisons had a significant correlation (p < .05), and each of the 13 markers correlated with at least one other marker. Parallel analysis was conducted to determine the maximum number of dimensions to be extracted from the model. Parallel analysis compares the sample eigenvalues (the amount of variance carried in each component) of the markers with the 95^th^ percentile of eigenvalues across multiple randomly generated datasets with same number of uncorrelated factors. Components with sample eigenvalues being higher than those of the randomly generated data can be extracted. Parallel analysis results revealed that the 13 markers were best reduced to three components, as determined by the sample eigenvalue being higher than that of the randomly generated data at up to three dimensions. This model would account for approximately 68% of the variance in the data (**Supp. Figure 3A**). Most of the biological measures were appropriately categorized into at least one component, as demonstrated by a factor loading of > 0.3^15,16^ (**Supp. Figure 3B**).

Notably, this initial model using all thirteen significant biological measures included four markers that did not have a strong loading into any of the three components (FSH in the blood and pituitary, and progesterone in the blood and hypothalamus). This may indicate that the dimension reduction by PCA may be being muddied by extraneous variables that are not immediately relevant to the model. Additionally, the grouping of biological measures into three components did not meet our needs in order to assign each sample a single composite score. Ignoring all but the first component would not have been appropriate, as the first dimension only accounted for approximately 38% of the variance within the dataset, and some biological measures with strong relevance to menopausal status had a low factor loading with the first component (e.g. AMH and FSH in the blood). Thus, we sought to perform a new analysis including only markers with established relevance to menopausal status (AMH, FSH, estradiol, and progesterone within the blood), and markers that were correlated with the aforementioned established biologically relevant measures (FSH and *FSH* gene expression in the pituitary and estradiol within the hypothalamus) (**Figures 2 and 4; Supp. Figure 2**). This led to a final model with seven biological markers for characterizing menopausal status. Of the 21 comparisons made between these seven markers, 15 comparisons had a significant correlation (p < .05), and every individual marker correlated with at least three others (**Figure 5**). This suggested a strong model with highly correlated factors. This suggestion was supported by parallel analysis, which revealed that the data was best reduced to a single dimension that accounted for approximately 47% of the variance within the dataset (**Figure 6A**). All markers had factor loading > 0.3 in magnitude, indicating that each factor had a moderate correlation with the final component score (**Figure 6B**).

**Figure 5:**
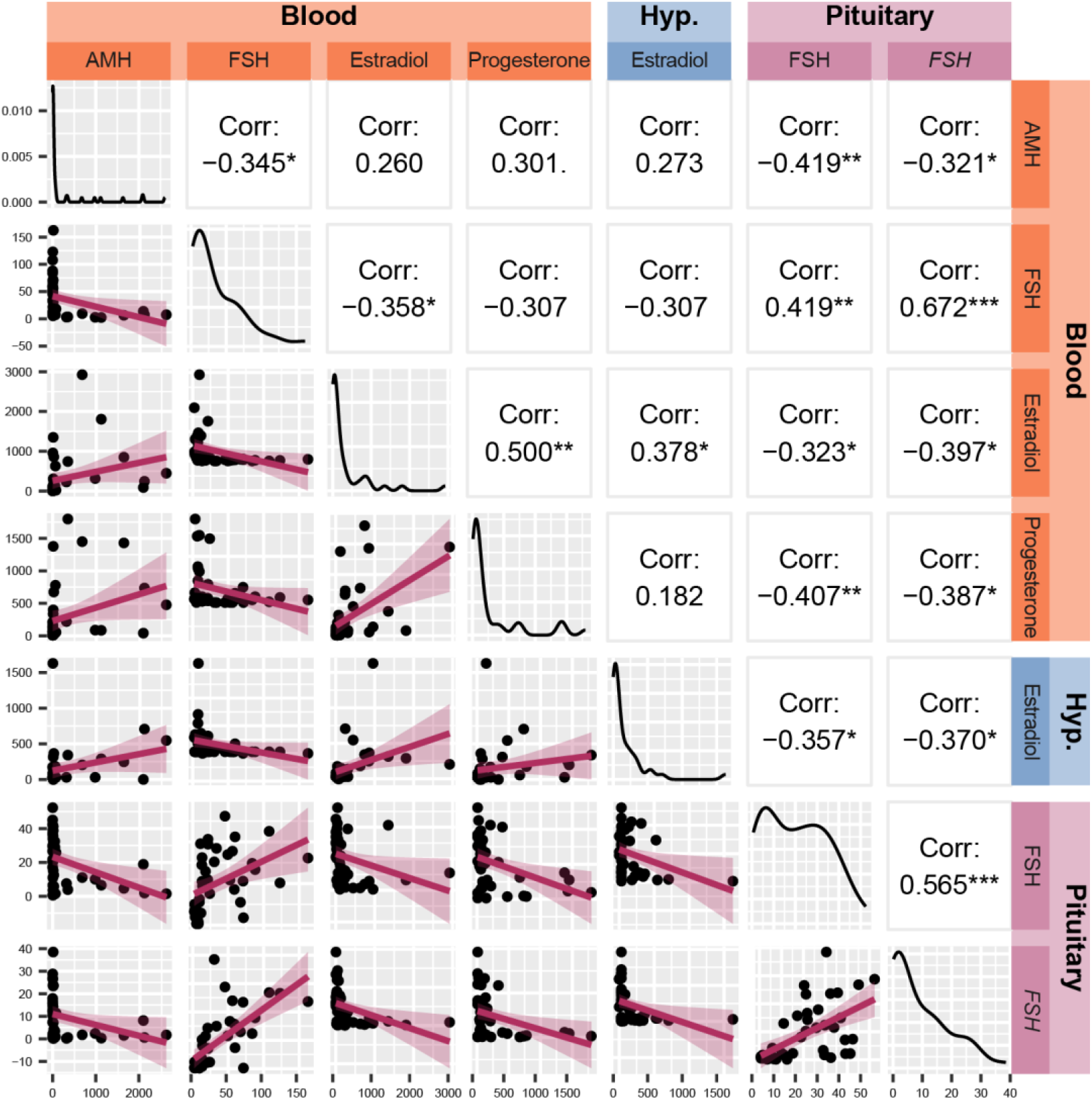
Correlation of seven biological measurements included in principal component analysis. The top-left/bottom-right diagonal represents distribution of each measurement, demonstrating a non-normal distribution for all seven markers. Below the diagonal are scatterplots of each pair of markers with the linear regression lines imposed. Above the diagonal are Pearson’s correlation coefficients for each marker with each other.*p < 0.05, **p < 0.01, ***p < 0.001 Glossary: AMH – Anti-Müllerian hormone; FSH – Follicle-stimulating hormone (protein); *FSH* – Follicle-stimulating hormone (gene).

**Figure 6:**
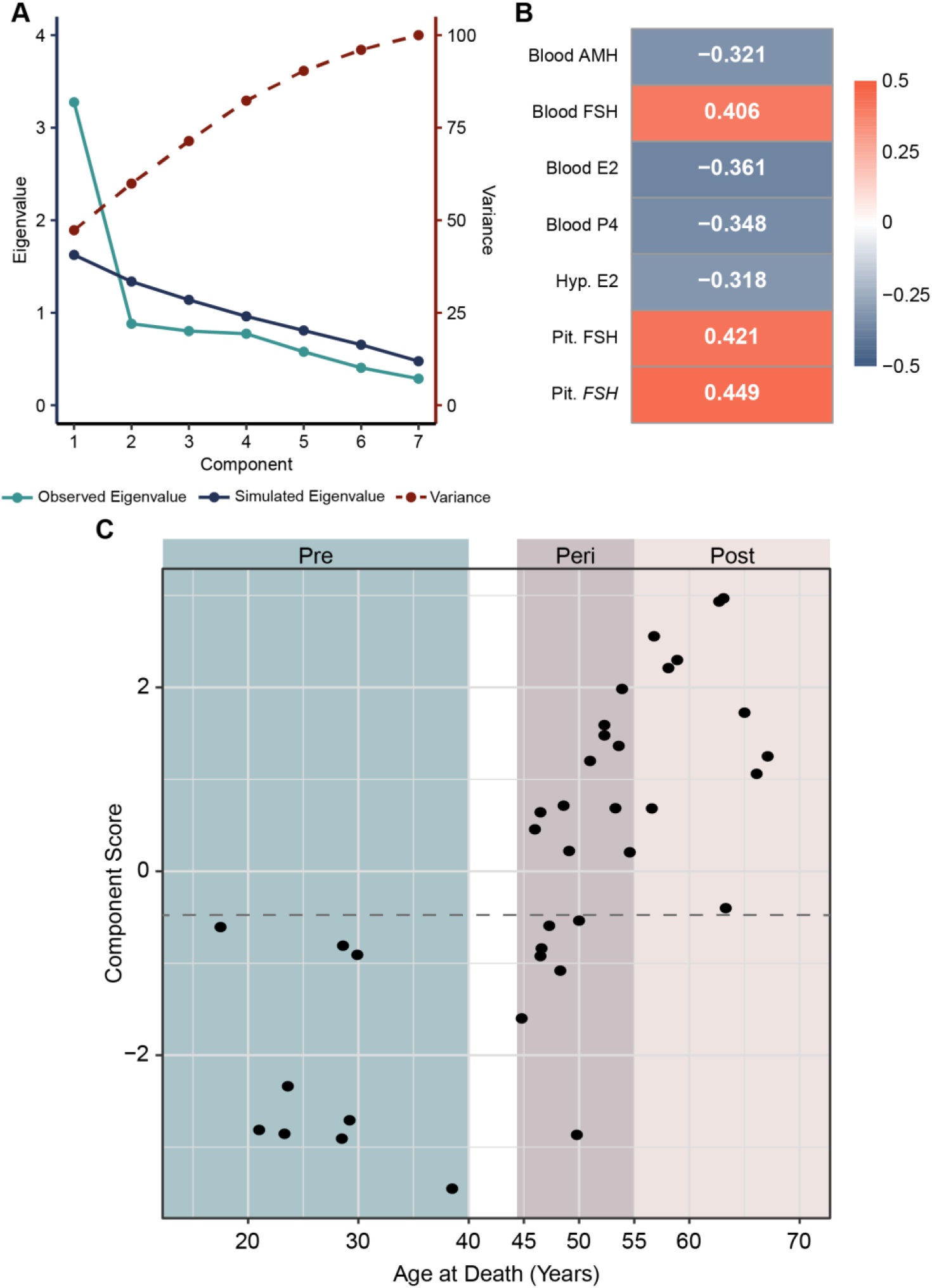
Characterization of samples based on a composite measure calculated by principal component analysis. **A)** The seven selected biological markers are most appropriately combined into a single component, as represented by the observed Eigenvalue (light blue line) being higher than the simulated Eigenvalue (dark blue line) only at the first component. This single component accounts for approximately 47% of variance in the data (dashed red line). **B)** All seven biological measures had at least moderate correlation with the final component score, as demonstrated by the absolute value of factor loadings falling between 0.3-0.5. A positive factor loading indicates that as this measure increases the component score increases, and a negative factor loading indicates that as this measure decreases the component score increases. **C)** Plot of chronological age vs. component score. All samples in the premenopause group had a component score of < -0.6, and all samples within the postmenopause group had a component score of > -0.4. A cutoff value to classify samples in the perimenopause group was set halfway between these two limits at -0.5 (dashed line). Glossary: AMH - AMH – Anti-Müllerian hormone; E2 – Estradiol; FSH – Follicle-stimulating hormone (protein); *FSH* – Follicle-stimulating hormone (gene); P4 – Progesterone.

Since our cohort included the individuals with psychiatric diagnoses (**Table 2**), we next wanted to examine whether having a psychiatric diagnosis impacts the final composite measure. Psychiatric diagnoses were characterized as binary variables (history of psychiatric diagnosis is either present or not present), and multiple factor analysis^17^, which extends PCA to handle binary variables, was performed to determine if psychiatric diagnosis would significantly impact categorization of menopausal status. Two sets of multiple factor analyses were performed: one collapsing all diagnoses into a single binary variable (using Axis 1 only), and one separating each diagnosis out into three separate binary variables. None of the psychiatric diagnosis groupings had strong associations with the component of the seven biological measurements. Thus, all psychiatric diagnoses were excluded from the final composite score calculation.

Finally, the component score was calculated for each sample in order to categorize them into one of two groups – “pre-menopausal like” or “post-menopausal like” groups. Calculation of the component score revealed that the premenopausal and postmenopausal samples created two distinct groups, with every premenopausal sample having a component score < -0.6 and every postmenopausal sample having a component score > -0.4 (**Figure 6C**). Thus, based on the measurements obtained in our current cohort, the cutoff value of -0.5 can be used to categorize the 18 peri-menopausal samples into two groups– pre- or post-menopausal like groups (**Supp. Table 1**). Three of the 21 perimenopausal samples could not have a component score calculated due to missing at least one of the seven biological measures included in the model.

After calculating the composite measure, we wished to determine whether this model could more reliably characterize each sample than manual characterization based on measures used in clinical settings. An experimenter blinded to each sample’s age and composite measure examined AMH and FSH levels within each of the 42 samples and characterized each measurement as “premenopausal levels” or “postmenopausal levels” based on proposed clinical standards (postmenopausal levels defined as AMH < 10 pg/mL, FSH > 25 mIU/mL)^2,3^ (**Supp. Table 1**). For all 10 premenopausal samples, grouping by chronological age, manual characterization based on AMH and FSH, and composite score all agreed on group assignment. For the remaining 32 perimenopausal and postmenopausal samples, 10 samples had conflicting AMH and FSH measurements that would make it difficult to characterize menopausal status. Of these 10 samples, three were older than 58 years old, indicating that these two biological measures alone may be unreliable even for individuals with a high confidence of postmenopausal status due to chronological age. All three samples were notably correctly characterized as postmenopausal by the composite score despite conflicting measurements in AMH and FSH. Of the seven perimenopausal samples with conflicting AMH and FSH measurements, six had composite scores well outside the cutoff range of -0.6 to -0.4, indicating that the other five measures that contributed to the model provided stronger confidence in the characterization of these samples. One sample had a composite score of -0.59, which falls just within the cutoff range. However, this sample came from an individual with unspecified infertility indicated in their history, which could lead to a low AMH measurement raising the component score despite all of the other six biological measures aligning with premenopausal status. All this together supports this composite measure as a more reliable way to characterize individuals within the perimenopausal age range when proposed clinical measures may conflict with each other.

### Alternative, tissue-specific component scores

Finally, considering that many brain banks do not have blood available and that working with multiple tissues may not be feasible or cost effective for researchers, we calculated tissue-specific component scores for hypothalamic markers, pituitary markers, and blood markers. We first selected tissue-specific markers that were among the 13 significant markers we initially revealed (**Table 2**). For the hypothalamus, these markers included estrone, estradiol, progesterone and DHEA (**Supp. Figures 4A, 5A-B**). Unlike the model using seven selected biological markers as described previously, the hypothalamus-only PCA did not generate a model with a clear cutoff between premenopausal and postmenopausal groups, making characterization of the perimenopausal samples less definitive. All but one premenopausal sample had hypothalamus composite measures of > -0.3, and all but two postmenopausal samples had hypothalamus composite measures of < -0.7, meaning that the most appropriate cutoff score would be between these two values (**Figure 7A**). With a cutoff of -0.5, 32 out of 37 samples (86.5%) agreed between the hypothalamus component score and the 7-marker component score. There was a moderate, negative correlation between these two scores (r = -0.54, p < 0.001; **Figure 7B**).

**Figure 7:**
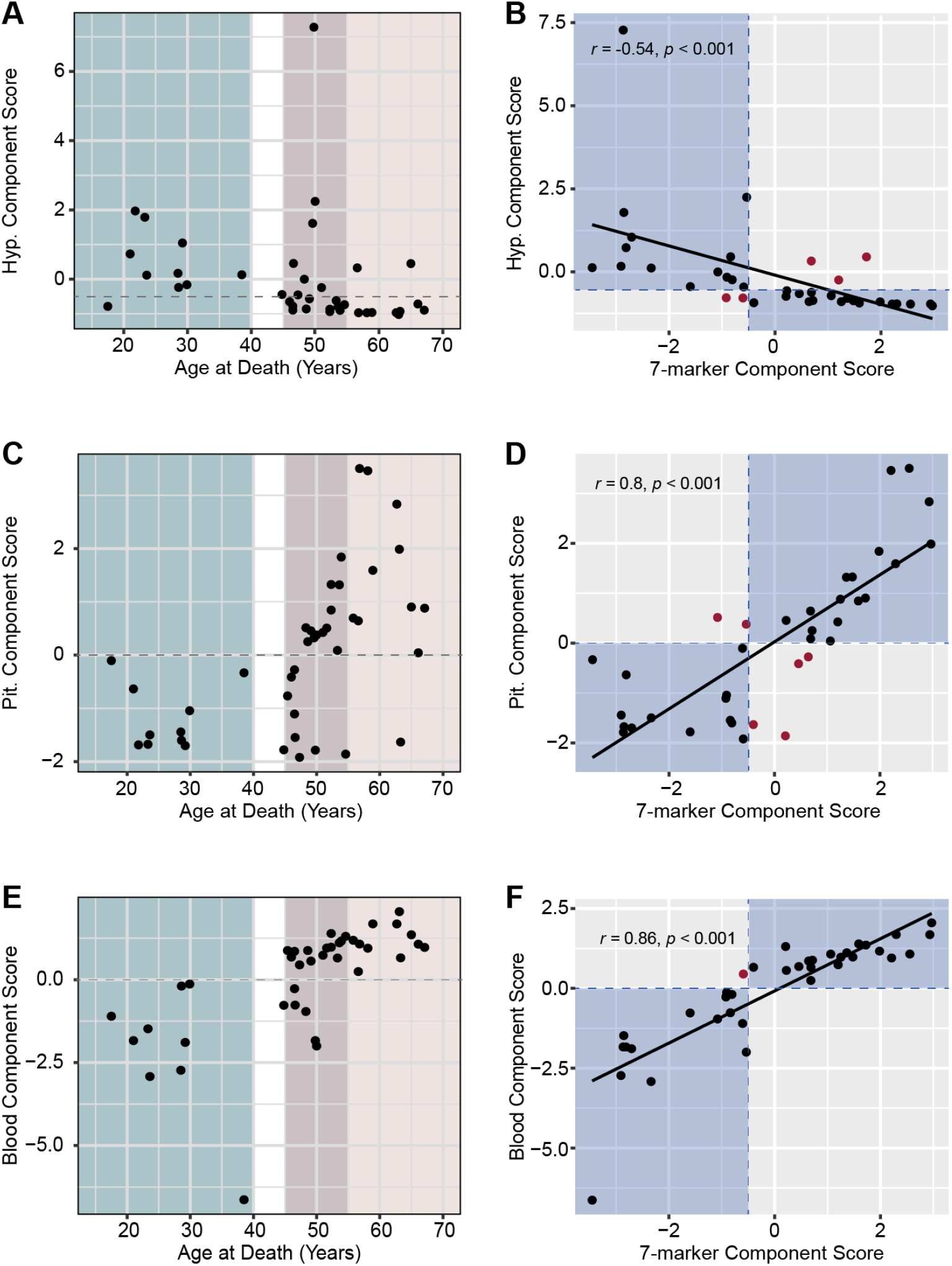
Tissue-specific component scores. **A)** Plot of chronological age vs. hypothalamus component score. 9/10 premenopausal samples had a component score of > -0.3, and 10/11 postmenopausal samples had a component score of < -0.7. A cutoff value to classify samples in the perimenopause group was set halfway between these two limits at -0.5 (dashed line) **B)** The hypothalamus component score moderately correlated with the 7-marker component score. A total of 5 samples disagree between hypothalamus and 7-marker scores. **C)** Plot of chronological age vs. pituitary component score. All samples in the premenopause group had a component score of < 0, and all samples within the postmenopause group had a component score of > 0. **D)** The pituitary component score strongly correlated with the 7-marker component score. A total of 6 samples disagree between pituitary and 7-marker scores. **E)** Plot of chronological age vs. blood component score. All samples in the premenopause group had a component score of < 0, and all samples within the postmenopause group had a component score of > 0. **F)** The blood component score strongly correlated with the 7-marker component score. Only one sample disagreed between blood and 7-marker scores. On correlation plots **(B, D, F)**, blue areas indicate where respective tissue-specific score classification aligned with 7-marker composite measure classification, and red dots indicate samples with disagreement between the two classifications.

For the pituitary component score, we combined pituitary FSH protein levels and gene expression for *FSH* and *GNRHR* (**Supp. Figures 4B, 5C-D**). Importantly, the pituitary score had a clear distinction between the premenopausal and postmenopausal groups in their component score, with all premenopausal samples having a component score < 0 and all postmenopausal samples having a component score > 0 (**Figure 7C**). With this cutoff, 31 out of 37 (83.8%) samples agreed between the pituitary component score and the 7-marker component score. There was a strong positive correlation between these two scores (r = 0.80, p < 0.001) (**Figure 7D**).

Lastly, we calculated two blood component scores: one with all six markers found to have significant differences between the premenopausal and postmenopausal groups (AMH, FSH, estrone, estradiol, progesterone, and DHT) (**Supp. Figures 4C, 5E-F**), and one only containing the four steroid hormones (**Supp. Figures 4D, 5G-H**). Similar to the pituitary component score, the full blood component scores had a distinct separation between the premenopausal and postmenopausal groups, with all premenopausal samples having scores < 0 and all postmenopausal samples having scores > 0 (**Figure 7E**). With this cutoff, 36 out of 37 samples (97.3%) agreed between the blood component score and the 7-marker component score. There was a strong positive correlation between these two scores (r = 0.84, p < 0.001) (**Figure 7F**). Conversely, the removal of AMH and FSH measures in the steroids-only score removed the clear distinction between premenopausal and postmenopausal groups, with some overlap of component scores. All but one premenopausal sample had a steroids-only component score of > -0.2, and all postmenopausal samples had component scores < -0.2 (**Supp. Figure 6A**). With a cutoff of -0.2, 35 out of 37 samples (94.6%) agreed between the steroids-only component score and the 7-marker component score. Unlike the full blood component score, the steroids-only score had a strong negative correlation with the 7-marker score (r = -0.73, p < 0.001) (**Supp. Figure 6B**). These analyses reveal that future application of this approach in classifying premenopausal-like and postmenopausal-like cases is feasible in cases of limited tissue availability, as all four tissue-specific models generally have high agreement with the full composite measure (**Supp. Table 2**). Finally, to examine the robustness of the PCA results, we performed an additional PCA using the Spearman’s rank correlations of the markers, as well as the partial Pearson’s correlations of the markers controlling for PMI, hypothalamus RNA integrity number (RIN), and pituitary RIN. The eigenvalues and the factor loadings of these results were consistent with those presented previously (**Supp. Figure 7**).

## Discussion

With the increasing life expectancy and the average age of menopause being 51 years, it is expected that women and other people with ovaries will spend nearly half of their lives in menopause. Importantly, although the MT is associated with the increased risk for many brain disorders^6,8,10,11^, the underlying cellular and molecular mechanisms remain poorly understood. To enable postmortem investigations of the human brain across the MT, here we reveal the biomarkers and composite measures across three different tissues that can be used to characterize an individual’s menopausal status postmortem.

Twelve months of amenorrhea are currently required for a formal diagnosis of menopause in living humans, but this is an arbitrary, agreed-upon clinical requirement that is not clearly defined by any biological marker. Revealing the endocrinology underlying the uterine bleeding that characterizes the MT has been challenging because of the marked variability in hormones during this period. Previous studies have proposed using serum levels of FSH^5,14^ or AMH^3,18^ to either diagnose or predict menopause, although no standardized measures have been established so far. While studies in living humans at least provided some guidelines for reference biological measures and the menstrual cycling pattern can be directly inquired from patients and study participants, there is practically no information on how to determine reproductive status from postmortem human tissues as this individual or clinical information is typically lacking in tissue repositories.

We chose three tissues that synthesize or carry HPG-related hormones of direct relevance to reproductive function: blood, the hypothalamus, and pituitary gland. While plasma or serum are typically tissues of choice for exploring the reproductive status in living humans, it is important to note that brain tissue repositories either do not carry blood/serum or may only provide frozen whole blood samples. Importantly, here we show that AMH, FSH, and steroid hormones can be reliably detected in frozen postmortem blood samples, corresponding well to the values observed in clinical serum samples. As expected, blood AMH, FSH, estradiol, and progesterone are among the strongest menopausal markers. Similar to findings from clinical populations, AMH or FSH by themselves are not sufficient to clearly determine reproductive state postmortem. Rather, our findings show that using the postmortem blood composite measure of the six strongest blood biomarkers is allowing for high confidence prediction of whether the sample comes from a “pre-menopausal like” or “post-menopausal like” individual. If measuring all six markers is cost prohibitive since they require three separate tests, the composite measure of four steroid hormones in blood also allows for a fairly good prediction of the menopausal status.

Another tissue of interest was the hypothalamus, because this region is both involved in the HPG axis and has high density of ovarian hormone receptors^19^. In addition, for brain banks that do not provide access to blood, the hypothalamus can provide an alternative tissue source to determine steroid hormone levels. Previous studies showed that circulating estradiol^20^ and progesterone^21^ levels are reflected in the hypothalamus and that they differ between postmenopausal and premenopausal women. However, in a small sample size study (N=11), it was shown that estradiol levels are significantly reduced during postmenopause in the preoptic area but not in the medial and basal hypothalamus^20^ as compared to premenopause. In addition, this study did not find a significant correlation between serum and brain concentration of estradiol. However, here we show a significant correlation between steroid hormone levels in blood and in (an unspecified area of) the hypothalamus, which may reflect a higher sensitivity of the method that we used compared to the previous study (HPLC-Mass Spectrometry vs. Radioimmunoassay).

One important consideration regarding the hypothalamus is that this is a highly heterogeneous region and lack of consistent brain dissections may lead to difficulty in finding appropriate biomarkers from this brain region. Indeed, with the acquired material from the hypothalamus, we were not able to find any MT-related changes in hypothalamic gene expression, which is likely due to our inability to acquire a more specific part of the hypothalamus. As an example, previous studies have found either increased^22^ or decreased^23^ expression of the kisspeptin (*KISS1*) gene in the human hypothalamus following menopause, depending on a specific hypothalamic region. Here, we did not observe a difference in *KISS1* expression across the groups and individual values demonstrated extreme variability consistent with its region-specific expression. For this reason, our finding that steroid hormone levels are correlated with their blood levels reveal the possible utility of hypothalamic steroid measurements in the investigations of the menopausal effects on the brain, as this effect may be less region-specific than gene expression. In addition, there is a modest correlation between multi-tissue and hypothalamic-specific composite measures for the menopausal status implying that the hypothalamus can represent an alternative tissue source for studies of the MT and menopause.

Among the most remarkable and less expected findings was the utility of the pituitary gland as an alternative tissue source for determining the menopausal status. While FSH is produced in the pituitary, the FSH assay is typically performed using serum, not the pituitary. Here, however, we show that pituitary FSH represents an excellent menopausal marker, at both protein and gene expression levels. Our data are consistent with previous findings that FSH levels in serum are primarily regulated at the level of FSH production rather than through regulation of FSH release^24^. But, importantly, combining pituitary FSH protein and gene expression with the pituitary *GNRHR* gene expression gives almost a perfect composite score that resembles the results of applying multi-tissue composite measure.

What is clear from this study is that having multiple, highly-correlated biomarkers increases the ability to predict an individual’s menopausal status. While combining all three tissue indices together gives the most precise information, having markers from blood or pituitary only provide highly reliable results as well. Finally, while not ideal, the hypothalamus can also provide a solid, alternative tissue source if blood or pituitary are not available to researchers. The results of our study provide a long missing method to determine the reproductive status postmortem, allowing the molecular and cellular studies of the MT effect on the human brain. By enabling the study of mechanisms underlying increased neuropsychiatric risk during the MT, we open the path for the hormone status-informed, precision medicine approach in women’s mental health.

## Acknowledgements

This work was supported, in part, by the National Institutes of Mental Health (NIMH) under Award Number R01MH123523 and a Fordham University’s Faculty Research Grant (to M.K.) The Human Brain Collection Core (HBCC) is supported by the NIMH-IRP project ZIC MH002903. We would like to thank Brian Nofsinger for his assistance with steroid hormone quantification.

## Author contributions

M.K. conceived and directed the project and provided funding; M.T., H.C., P.A., S.M., and M.K. designed the study; M.T. prepared samples for the analyses and performed ELISA and gene expression experiments; M.T. and H.C. performed data analysis; A.G. performed manual characterization of the samples; P.A. and S.M. provided posmortem tissue specimens, M.T., H.C., and M.K. interpreted the data and constructed the figures; M.T. and M.K. wrote the article with input from all co-authors. All authors approved the final version of the manuscript.

## Materials and Methods

### Subjects

The HBCC collected brains, surrounding tissues and blood primarily from medical examiners in Virginia and Washington, D.C. with the permission from the next-of-kin. A telephone interview was conducted with the next-of-kin to gather demographic information and medical history. This was complemented by collection of medical records where available. Medical history was reviewed, and a consensus clinical diagnosis was reached by two psychiatrists based on DSM-IV or DSM-5 criteria. Postmortem brains were obtained from both individuals with no neuropsychiatric illness in their lifetime and individuals with psychiatric disorders (including mood, anxiety, and substance use disorders; **Table 1**). All cases were assessed by a neuropathologist and found to be free of neurodegenerative disease.

Samples from a total of 42 individuals were received from the HBCC. Each sample was categorized based on age at the time of death as either “premenopausal” (< 40 years old), “perimenopausal” (45-55 years old), or “postmenopausal” (> 55 years old). For every sample, we received ∼200μg of tissue from the hypothalamus, ∼500μg of tissue from the pituitary gland including both anterior and posterior portions, and 500-1000μL of whole blood.

### Sample preparation

Brain collection and processing was performed as previously described^25^. Briefly, brains were sectioned into ∼1cm coronal slabs, and slabs were flash frozen in a slurry of isopentane and dry ice prior to storage at - 80ºC. Upon receipt of tissue request, slabs were removed from storage and blocks of tissue were dissected, maintaining frozen temperature throughout processing. The HBCC provided tissue from the hypothalamus and pituitary as well as aliquots of whole blood for analysis.

Tissue blocks received from the HBCC were pulverized to a coarse powder over dry ice. Pulverized tissue samples were separated into aliquots for separate analyses. Approximately 100mg of hypothalamic tissue was set aside for steroid analysis, and 50-100mg of hypothalamic tissue was used for RNA isolation. Between 25-75mg of pulverized pituitary tissue was used each for RNA isolation and protein extraction. Pulverized tissue was stored at -80ºC until further processing.

Whole blood was thawed and centrifuged for 20 minutes at 1500xg at 4ºC to separate coagulation before aliquoting. 200µL of whole blood was set aside for steroid analysis and two 100μL aliquots were set aside for ELISA assays. All aliquots were stored at -80ºC until analysis.

Three premenopausal samples also had serum provided, which were used during assay optimization to confirm whether each assay was compatible with whole blood and determine the ratio of whole blood measures to plasma measures of AMH and the 14 steroids. All measures had a similar pattern of whole blood measuring at around 50% of the values for plasma (**Supp. Table 3**).

### Steroid panel

Steroid panel concentration was quantified with high-performance liquid chromatography with tandem-mass spectrometry (HPLC-MS/MS) by OpAns, LLC (Durham, NC). The reference standards for aldosterone, androstenedione, corticosterone, cortisol, cortisone, dehydroepiandrosterone (DHEA), deoxycorticosterone (DOC), 11-deoxycortisol, dihydrotestosterone (DHT), estradiol, estrone, 17OH-progesterone, progesterone, and testosterone were prepared as individual stock solutions at 1 mg/mL in acetonitrile (ACN) and dimethyl sulfoxide (DMSO) (50:50), then further diluted in ACN and DMSO (50:50) to create calibration and quality control standards. Stable isotropic labeled internal standards for each analyte were added to calibration standards, quality control, and matrix samples.

Analytes and their internal standards were extracted from whole blood by liquid-liquid extraction using methyl tert-butyl ether (MTBE), then evaporated to dryness and reconstituted in MeOH:water. Hypothalamus samples were homogenized in MeOH and EtOAc (50:50), and the resulting homogenate was centrifuged and the supernatant evaporated. The extract was then reconstituted in MeOH and diluted with water, then loaded onto a conditioned Biotage Evolute Express ABN solid phase extraction cartridge and subsequently washed with water and MeOH. The sample was then eluted with MeOH and evaporated to dryness, then once more reconstituted in MeOH:water. The reconstituted blood and hypothalamus extracts were then analyzed with HPLC-MS/MS using a 1290 series HPLC system (Agilent Technologies) and Agilent 6495 series triple quadrupole tandem mass spectrometer. HPLC-MS/MS data were acquired and quantified using the software application MassHunter Workstation Data Acquisition for Triple Quad version 10.1/ build 10.1.67 (Agilent Technologies).

### Pituitary protein extraction

An extraction buffer of phosphate buffered saline (pH 7.4) (PBS) containing 1X protease/phosphatase inhibitor cocktail was prepared and kept on ice during use. Pulverized pituitary was added to 1mL of chilled extraction buffer and homogenized, then centrifuged for 5 minutes at 5000xg at 4ºC. The supernatant was then removed and placed into a new chilled tube on ice and the tissue pellet was discarded. Total protein concentration was quantified with a Pierce BCA protein assay kit (Thermo Scientific) according to manufacturer instructions. Protein extract was stored at -80ºC until analysis.

### FSH ELISA

Whole blood aliquots and pituitary protein extracts were thawed over ice, and FSH concentration was quantified using an ELISA kit (Novus Biologicals) according to manufacturer instructions. Whole blood was diluted to a 1:10 concentration in PBS, and pituitary protein extracts were sequentially diluted to 1:100 concentration twice in PBS for a final concentration of 1:10000. Diluted samples and prepared standards were loaded in duplicate into the pre-coated plates, then antibody solution was added. Plates were incubated at 37ºC for 1 hour, then washed three times with wash buffer. Following washing, HRP-conjugate was added and the plate was incubated at 37ºC for 30 minutes. The plates were washed once more, and development substrates were added before one final incubation at 37ºC for 15 minutes. Finally, stop solution was added and the plates were read with a microplate reader (BMG Labtech) at 450nm. Optical densities of duplicates were averaged and the optical density of the 0 standard (blank) was subtracted from every value. Sample concentrations were calculated by plotting the log of sample optical density along the standard curve generated by the log of standard optical density against the log of standard concentrations, then appropriately multiplied by the dilution factor to determine the total concentration. Pituitary FSH concentration was then divided by total protein concentration to determine the concentration of FSH per mg of extracted protein.

Blood intra- and inter-assay CV’s were 5.6% and 2.2%, respectively. Pituitary intra- and inter-assay CV’s were 4.2% and 3.5%, respectively.

### AMH ELISA

Whole blood aliquots were thawed over ice, and AMH concentration was quantified using the picoAMH ELISA kit (Ansh Labs) according to manufacturer instructions. Microplates were loaded with assay buffer, calibrators, and controls. 20μL of each whole blood sample was loaded in duplicate, then 80μL of sample diluent was added to each sample well to bring each sample well to a final 1:5 concentration. Plates were then incubated at room temperature for three hours while on a shaker at ∼600rpm. Plates were then washed five times with wash buffer, and antibody-biotin conjugate was added before another incubation at room temperature for one hour while shaking. Plates were washed again, then loaded with streptavidin-enzyme conjugate before incubating again at room temperature for 30 minutes while shaking. Plates were washed one final time, then loaded with TMB solution and incubated at room temperature for 10 minutes while shaking. Stop solution was then added and the plates were read using a microplate reader at 450nm. Sample concentrations were calculated by plotting the log of sample optical density along the standard curve generated by the log of calibrator optical density against the log of calibrator concentrations, then appropriately multiplied by the dilution factor to determine the total concentration. Intra- and inter-assay CV’s were 4.1% and 2.6%, respectively. Calculated concentrations for Controls I and II provided within each picoAMH kit were within stated concentration ranges.

### RNA isolation and RT-qPCR

Pituitary and hypothalamus tissue was first processed with Trizol reagent (Thermo Scientific) and RNA was isolated from the aqueous layer using the Allprep DNA/RNA Mini kit (Qiagen). Pulverized tissue was homogenized in 1mL Trizol reagent, 200uL of chloroform was added and mixed by inversion, then samples incubated at room temperature for 10 minutes. Samples were then centrifuged at 4ºC for 15 minutes at 12,000xg. Following centrifugation, the aqueous phase was removed and loaded into the Allprep DNA spin column. From this point samples were processed according to manufacturer instructions, with the RW1 wash step separated into two 350μL washes before and after DNase (Qiagen) added to the spin column to ensure elimination of gDNA within the sample. Following RNA elution, total RNA concentration was determined using a Qubit RNA High Sensitivity kit and Qubit fluorometer (Invitrogen). RIN was determined with Bioanalyzer using an RNA 6000 Pico kit (Agilent). Isolated RNA was stored at -20ºC until conversion. 500ng of RNA was reverse transcribed to cDNA using SuperScript III First-Strand Synthesis System (Invitrogen), and stored at -20ºC until analysis. cDNA was analyzed for gene expression with quantitative PCR using a Quant Studio 3 real-time PCR system (Applied Biosystems). Forward and reverse primers for each gene analyzed are listed in **Supplementary Table 4**. Both hypothalamus and pituitary genes of interest were normalized against three housekeeping genes each using the previously described method^26^. The housekeeping genes used for the hypothalamus were *CYC1, UBE2D2*, and *EIF4A2*^*27*,*28*^. The housekeeping genes used for the pituitary were *GAPDH, TBP*, and *PSMC4*^*29*,*30*^. Gene expression was calculated relative to the premenopausal group.

## Statistics

### Individual markers

Demographic information (**Table 1**) including age, education level, PMI, and RIN values were analyzed with one-way ANOVA with Bonferroni-Hochberg corrections for multiple comparisons. A total of 39 biological measurements were taken for each of the 42 samples, including steroid hormones, glycoproteins, and gene expression (**Table 2**). For steroid measurements that were detectable but below the limit of quantitation, the value was indicated to be half of the lower limit of quantitation for the purposes of performing statistical analysis. None of the markers met the assumption for normal distribution (Shapiro-Wilk p < 0.05), so non-parametric tests were used for group comparisons. All 39 analytes were compared with Kruskal-Wallis test, with Dunn’s test for pairwise comparison. Measurements that had a significant difference between premenopausal and postmenopausal groups as determined by Dunn’s test posthoc were used for subsequent analyses. Results are reported as Median (First quartile – Third quartile) by group.

### Composite measure

The composite measure was determined using principal component analysis (PCA), which is a statistical method of dimension reduction used to group variables that highly associate with each other into a single variable known as a “component.” Markers’ distribution was first examined using kernel density plots, demonstrating the non-normal distribution of each of the markers. Scatterplots of each pair of markers with linear regression lines imposed showed that most markers were linearly associated. We repeated the kernel density plots and scatterplots with linear regression lines after the natural-log transformation of the markers. Results were consistent and it was decided that no data transformation was needed.

Parallel analysis^15,31^ was used to determine the dimensionality of the markers. Observed eigenvalues were compared to the 95^th^ percentile of eigenvalues across multiple randomly generated datasets with same number of random, uncorrelated markers (simulated eigenvalues). Components with sample eigenvalues being higher than those of the randomly generated data can be extracted. The number of dimensions with an observed eigenvalue greater than that of the simulated eigenvalue was used for PCA. For PCA, all markers were standardized (mean = 0, standard deviation = 1). After determining the principal component model, multiple factor analysis^17^ was performed with confident Axis I diagnoses (including depression, anxiety, and substance use disorders) together with the selected biological measurements. Multiple factor analysis extends PCA to handle binary variables. We compared the degrees of associations between the diagnoses and the component with those between the biological measurements and the component. Notably a component score could not be calculated for any samples missing at least one measurement; this led to exclusion of one premenopausal sample, three perimenopausal samples, and one postmenopausal sample from the 7-marker composite measure. To explore the robustness of the PCA results, we performed an additional PCA using the Spearman’s rank correlations of the markers, as well as the partial Pearson’s correlations of the markers controlling for PMI, hypothalamus RNA integrity number (RIN), and pituitary RIN.

All analyses were conducted in R version 4.3.2 (https://www.R-project.org)^32^. Kruskal-Wallis comparison across groups was performed with kruskal.test function in the stats package. Dunn’s test for pairwise comparison was performed with the dunnTest function in the FSA package^33^ (https://CRAN.R-project.org/package=FSA). The fa.parallel function in package psych was used for parallel analysis^33^. The prcomp function in package stats was used for principal component analysis. The MFA function in package FactoMineR was used for multiple factor analysis (<u>CRAN - Package FactoMineR (r-project.org)</u>). The partial.r function in package psych was used for partial Pearson’s correlations.

**Supplementary Figure 1:**
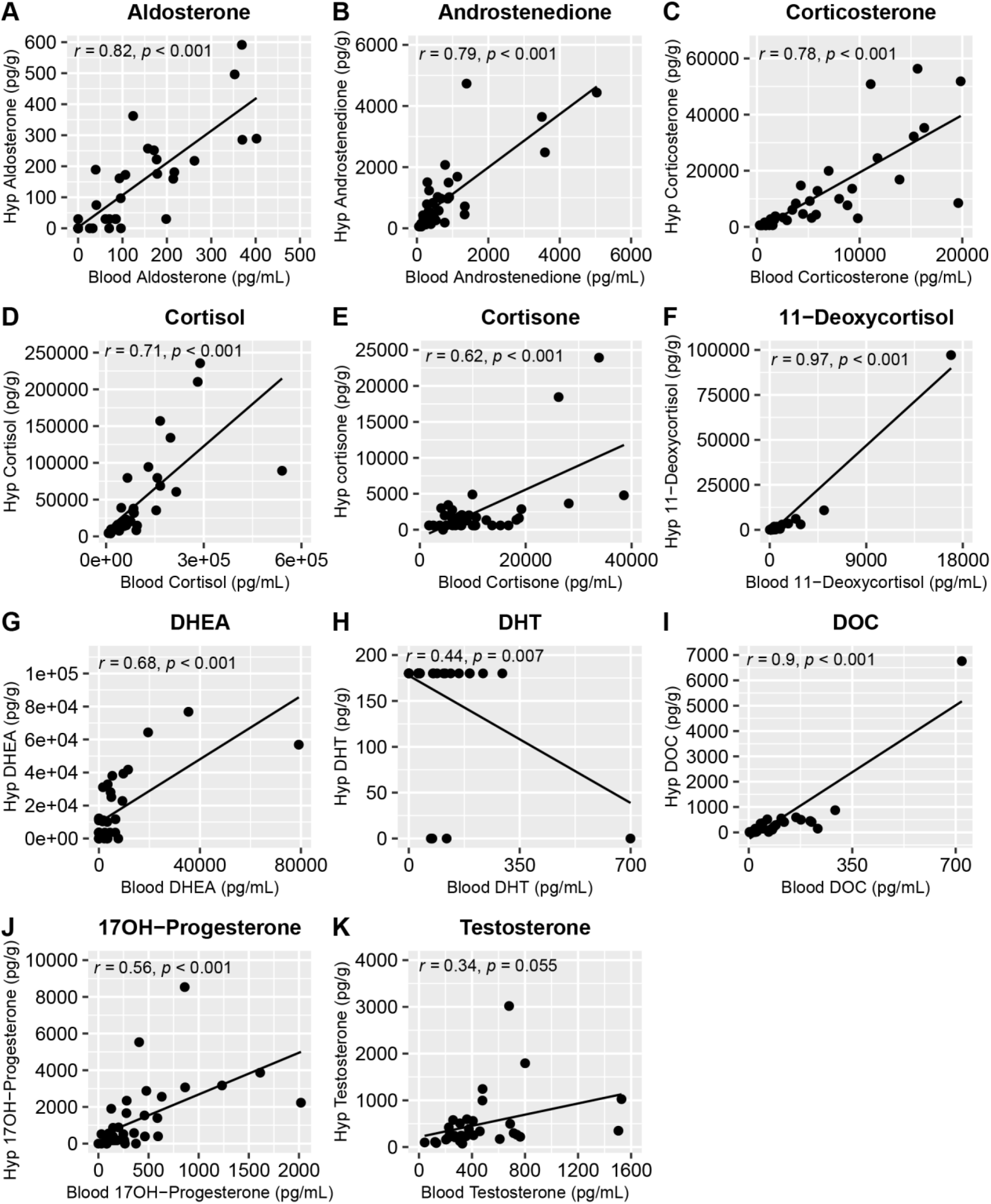
Correlation of blood vs. hypothalamic steroid measurements. All steroids, with the exception of DHT and testosterone, had significant moderate to very strong correlations between measurements from whole blood and measurements from hypothalamus tissue extract. Note that all hypothalamic DHT measurements were either undetectable (values replaced with 0 pg/g) or below the limit of quantitation (values replaced with ½ of the limit of quantitation). Glossary: DHEA – Dehydroepiandrosterone; DHT – Dihydrotestosterone; DOC – deoxycorticosterone.

**Supplementary Figure 2:**
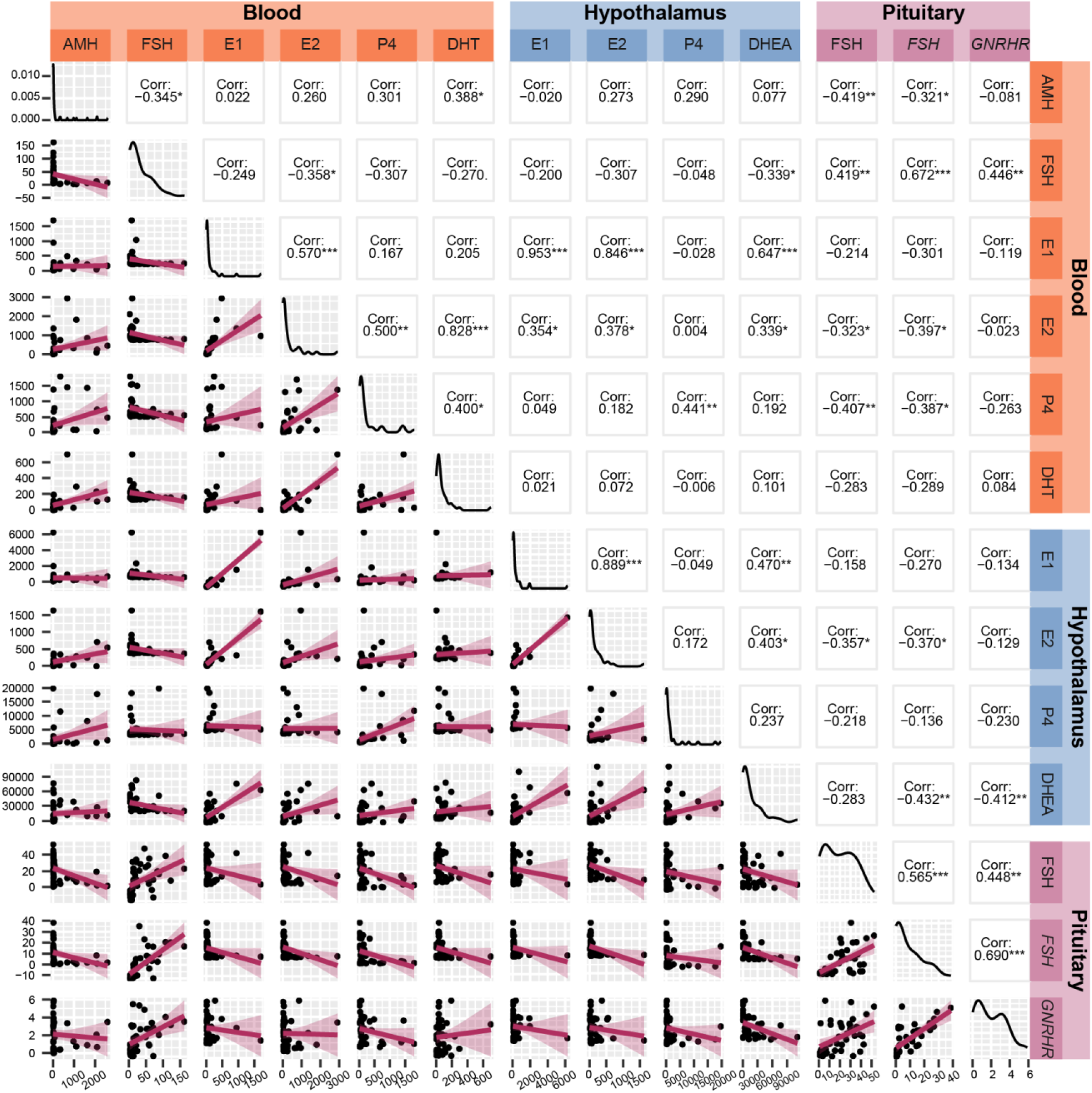
Correlation of thirteen biological measurements with significant differences between premenopausal and post-menopausal samples. The measurements were taken from blood (orange), the hypothalamus (blue), and the pituitary gland (purple). The top-left/bottom-right diagonal represents distribution of each measurement, demonstrating a non-normal distribution for all thirteen markers. Below the diagonal are scatterplots of each pair of markers with the linear regression lines imposed, showing that most markers are linearly associated. Above the diagonal are Pearson’s correlation coefficients for each marker with each other. *p < 0.05, **p < 0.01, ***p < 0.001. Glossary: AMH – Anti-Müllerian hormone; DHEA – Dehydroepiandrosterone; DHT – Dihydrotestosterone; E1 – Estrone; E2 – Estradiol; FSH – Follicle-stimulating hormone (protein); *FSH* – Follicle-stimulating hormone (gene); *GNRHR* – Gonadotropin-releasing hormone receptor (gene); P4 – Progesterone.

**Supplementary Figure 3:**
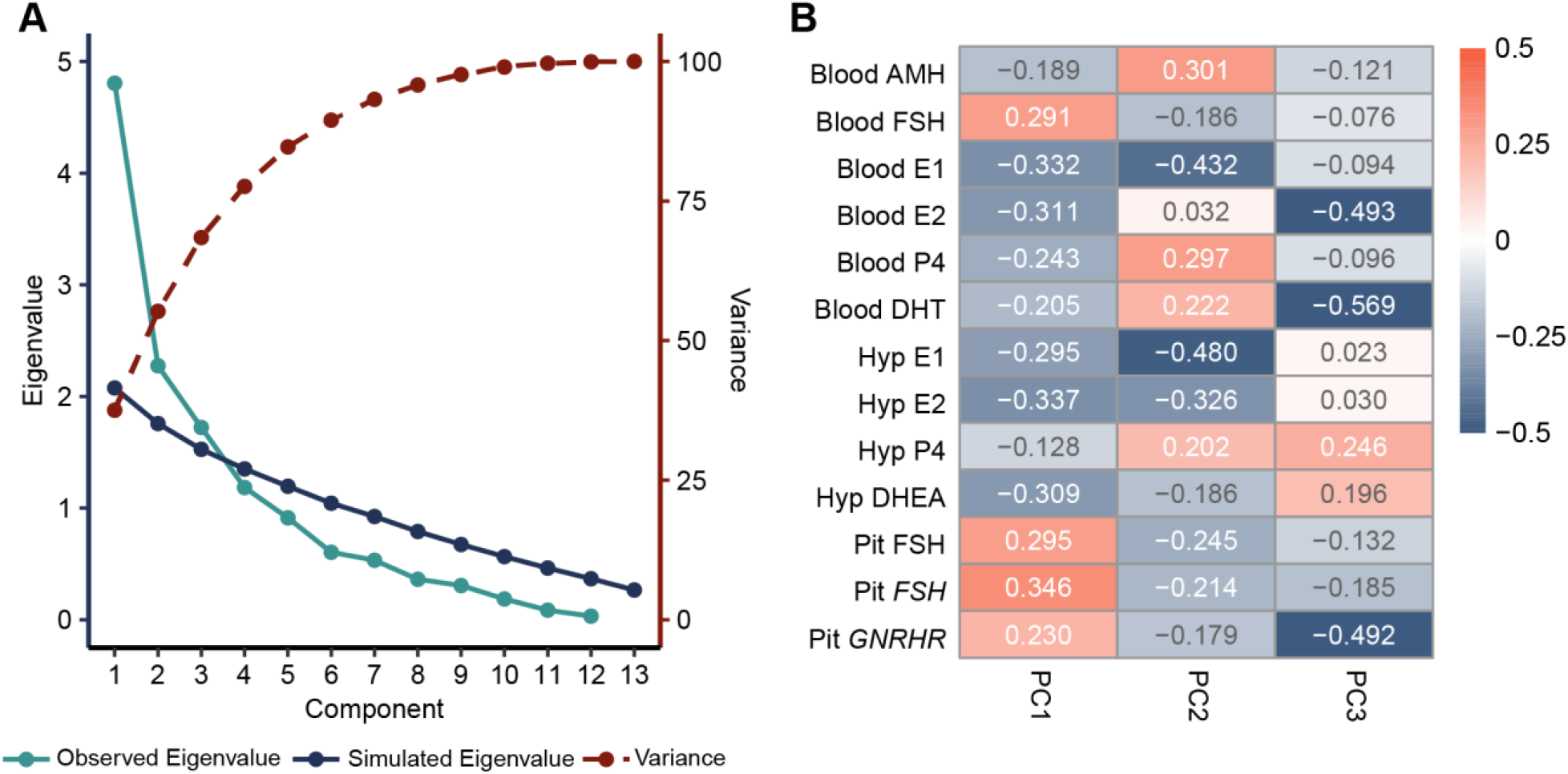
Principal component analysis for thirteen significant menopausal biomarkers. **A)** The thirteen significant biological markers are most appropriately combined into three components, as represented by the observed Eigenvalue (light blue line) being higher than the simulated Eigenvalue (dark blue line) with the highest variation in the dataset accounted for at three dimensions. Three dimensions accounts for approximately 70% of variance in the data (dashed red line). **B)** All thirteen significant biological measures have varying correlation with the three principal components. Glossary: AMH – Anti-Müllerian hormone; DHEA – Dehydroepiandrosterone; DHT – Dihydrotestosterone; E1 – Estrone; E2 – Estradiol; FSH – Follicle-stimulating hormone (protein); *FSH* – Follicle-stimulating hormone (gene); *GNRHR* – Gonadotropin-releasing hormone receptor (gene); P4 – Progesterone.

**Supplementary Figure 4:**
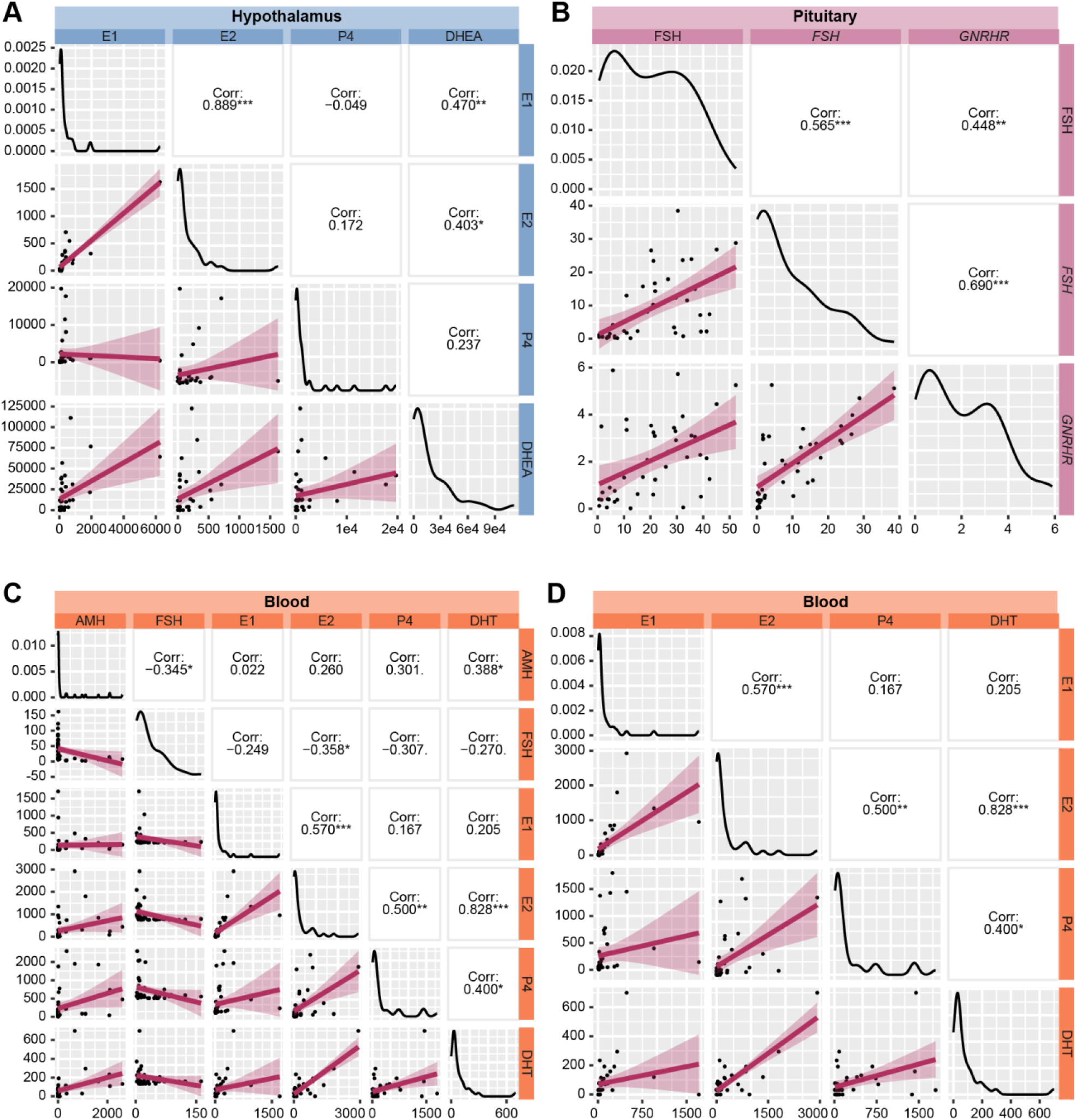
Correlations of biomarkers included in tissue-specific component score calculations. The top-left/bottom-right diagonal represents distribution of each measurement, below the diagonal are scatterplots of each pair of markers with the linear regression lines imposed, and above the diagonal are Pearson’s correlation coefficients for each marker with each other for **A)** hypothalamus, **B)** pituitary gland, **C)** blood, and **D)** steroid-only blood component scores. *p < 0.05, **p < 0.01, ***p < 0.001. Glossary: AMH – Anti-Müllerian hormone; DHEA – Dehydroepiandrosterone; DHT – Dihydrotestosterone; E1 – Estrone; E2 – Estradiol; FSH – Follicle-stimulating hormone (protein); *FSH* – Follicle-stimulating hormone (gene); *GNRHR* – Gonadotropin-releasing hormone receptor (gene); P4 – Progesterone.

**Supplementary Figure 5:**
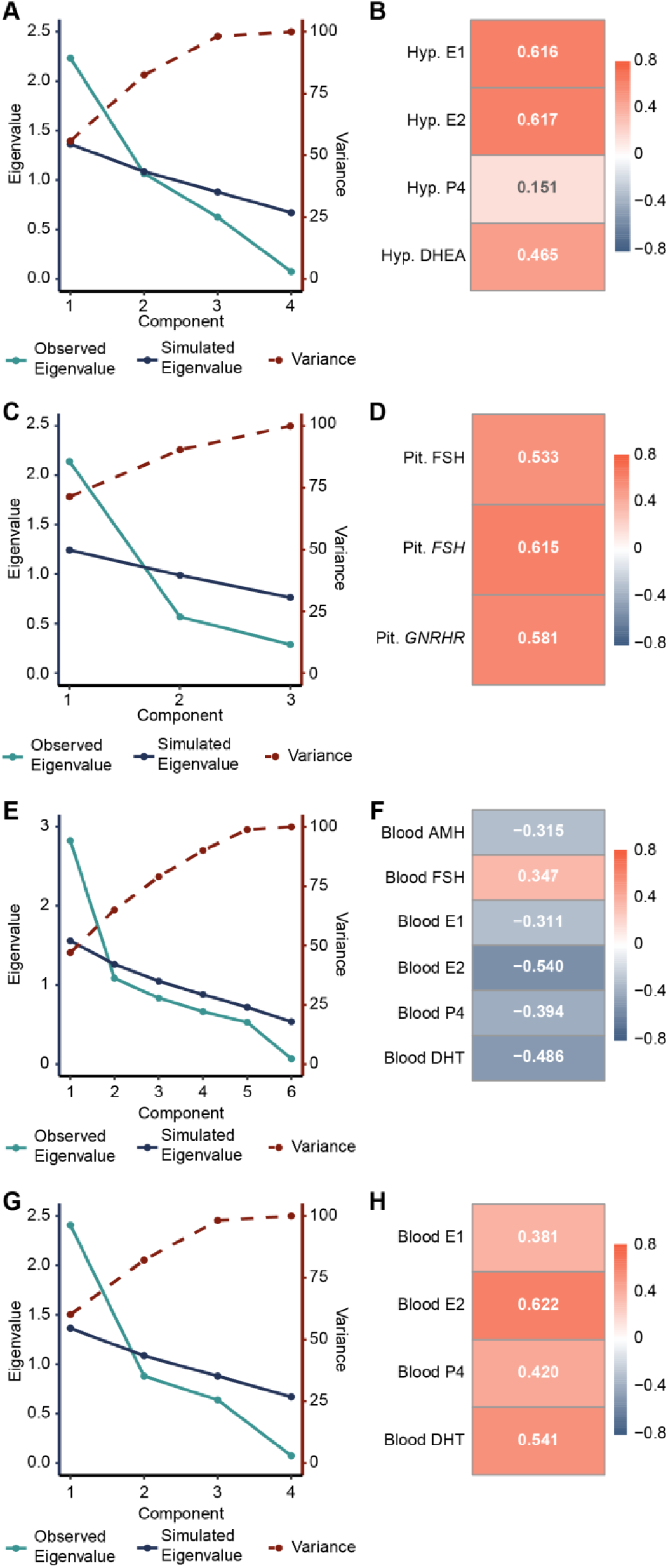
Principal component analysis for tissue-specific component scores. **A)** The hypothalamic markers are best combined into a single component, which accounts for approximately 56% of the variance in the data. **B)** All hypothalamic markers but progesterone loaded significantly into the component score. **C)** The pituitary markers are best combined into a single component, which accounts for approximately 71% of the variance in the dataset. **D)** All pituitary markers loaded significantly into the component score. **E)** The full blood markers are best combined into a single component, which accounts for approximately 47% of the variance in the dataset. **F)** All blood markers loaded significantly into the component score. **G)** The blood steroids are best combined into a single component, which accounts for approximately 60% of the variance in the data. **H)** All blood steroids loaded significantly into the component score. Glossary: AMH – Anti-Müllerian hormone; DHEA – Dehydroepiandrosterone; DHT – Dihydrotestosterone; E1 – Estrone; E2 – Estradiol; FSH – Follicle-stimulating hormone (protein); FSH – Follicle-stimulating hormone (gene); GNRHR – Gonadotropin-releasing hormone receptor (gene); P4 – Progesterone.

**Supplementary Figure 6:**
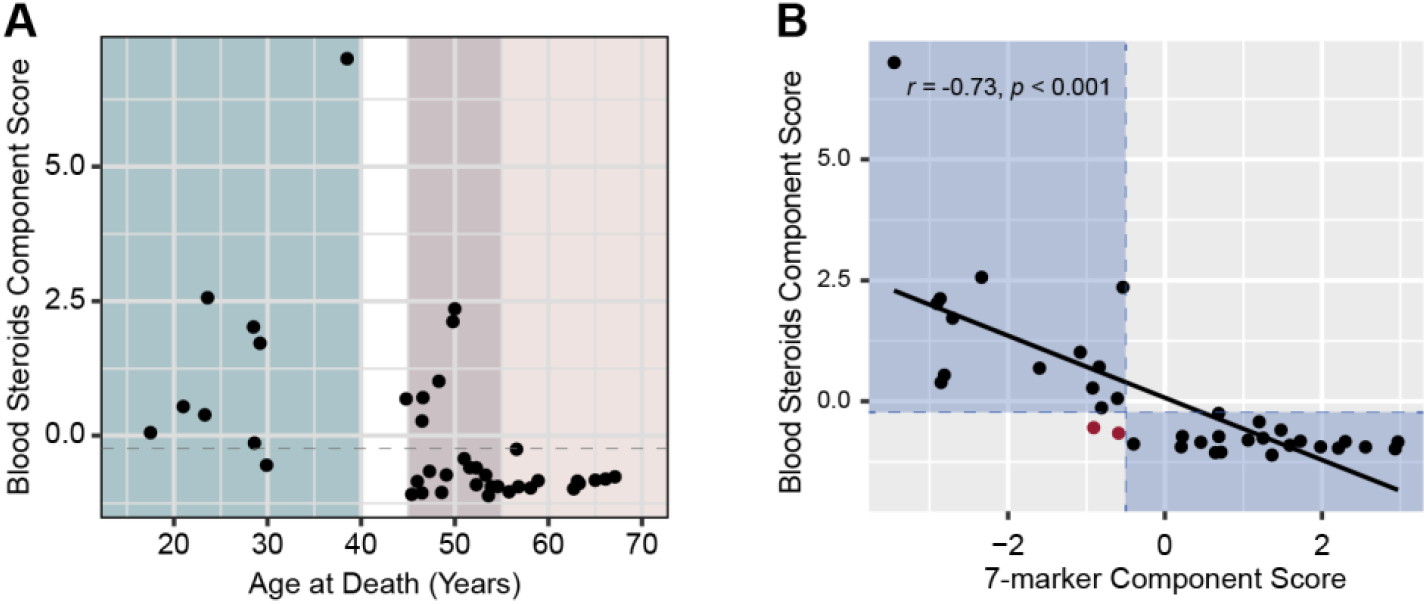
Characterization of steroids-only blood component score. **A)** There was some overlap in component score distribution between premenopause and postmenopause groups. With a cutoff of -0.2, all but one premenopause sample landed above the cutoff and all postmenopausal samples landed below the cutoff. **B)** Correlation between the blood steroids-only component score and the 7-marker component score was high, with only two samples having disagreement in classification between the two scores. Blue areas indicate where blood steroids-only tissue-specific score classification aligned with 7-marker composite measure classification, and red dots indicate samples with disagreement between the two classifications.

**Supplementary Figure 7:**
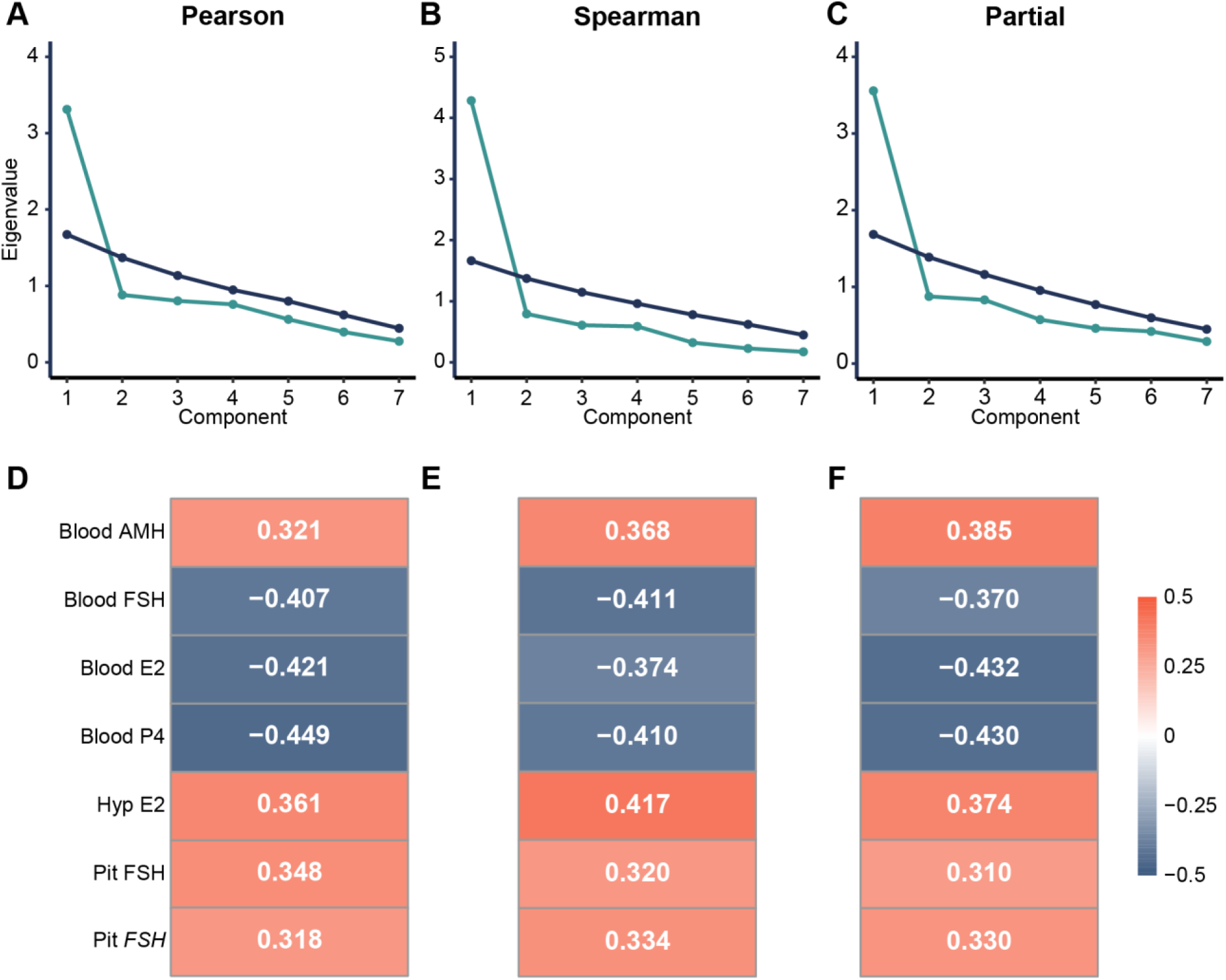
Supplemental PCA analyses with 7-marker model. **A-C)** Scree plots and **D-F)** factor loading scores for PCA models using Pearson, Spearman, and Pearson partial correlation accounting for PMI and RIN. All models have comparable factor loading scores to the final 7-marker model used for composite measure characterization of menopausal status. Glossary: AMH – Anti-Müllerian hormone; E2 – Estradiol; FSH– Follicle-stimulating hormone (protein); FSH – Follicle-stimulating hormone (gene); P4 – Progesterone.

